# ContamLD: Estimation of Ancient Nuclear DNA Contamination Using Breakdown of Linkage Disequilibrium

**DOI:** 10.1101/2020.02.06.938126

**Authors:** Nathan Nakatsuka, Éadaoin Harney, Swapan Mallick, Matthew Mah, Nick Patterson, David Reich

**Author notes:** co-first authors (Nathan Nakatsuka, Éadaoin Harney). Corresponding authors: Nathan Nakatsuka, Éadaoin Harney, and David Reich.

## Abstract

We report a method, *ContamLD*, for estimating autosomal ancient DNA (aDNA) contamination by measuring the breakdown of linkage disequilibrium in a sequenced individual due to the introduction of contaminant DNA, leveraging the idea that contaminants should have haplotypes uncorrelated to those of the studied individual. Using simulated data, we confirm that *ContamLD* accurately infers contamination rates with low standard errors (e.g. less than 1.5% standard error in cases with <10% contamination and data from at least 500,000 sequences covering SNPs). This method is optimized for application to aDNA, leveraging characteristic aDNA damage patterns to provide calibrated contamination estimates. Availability: https://github.com/nathan-nakatsuka/ContamLD.

## Background

Ancient DNA (aDNA) data has emerged as a powerful tool for learning about ancient population history, allowing direct study of the genomes of individuals who lived thousands of years in the past [1–3]. Unfortunately, these inferences can be distorted by contamination during the excavation and storage of skeletal material, as well as the intensive processing required to extract the DNA and convert it into a form that can be sequenced.

Accurate measurement of the proportion of contamination in ancient DNA data is important, because it can provide guidance about whether analysis should be restricted to sequences that show the characteristic pattern of C-to-T mismatch to the reference genome of authentic aDNA (if contamination is high) [4], or carried out at all. When analysis is restricted to focus only on sequences showing evidence of characteristic ancient DNA damage, the substantial majority of authentic sequences are usually removed from the analysis dataset, as only a fraction of genuinely ancient sequences typically carry characteristic damage. In addition, if a sample is contaminated by another individual with damaged DNA—which can arise for example as a result of cross-contamination from other specimens handled in the same ancient DNA laboratory—it is impossible to distinguish authentic sequences from contaminating ones based on the presence or absence of characteristic ancient DNA damage.

Current methods for estimating contamination have significant limitations. Methods based on testing for heterogeneity in mitochondrial DNA sequences (which are almost always homogeneous in an uncontaminated individual) can be biased, because there are several orders of magnitude of variation in the ratio of the mitochondrial to nuclear DNA copy number across samples. Thus, samples that have evidence of mitochondrial contamination can be nearly uncontaminated in their nuclear DNA, while samples that have no evidence of mitochondrial contamination can have high nuclear contamination [5]. Another reliable set of methods for estimating rates of contamination in ancient DNA leverage polymorphism on the X chromosome in males, including the popular *ANGSD* method [6–9] and an improved methodology that enhances power for low-coverage samples [10]. However, these methods do not work in females.

Several methods for estimating contamination rates in nuclear DNA from modern genomes have been published, including *ContEst* [11] and *ContaminationDetection* [12]. However, these methods generally rely on access to uncontaminated genotype data from the individual of interest or access to all possible contaminating individuals, neither of which is typically available for aDNA. Another method estimated modern human autosomal contamination in aDNA from archaic Denisovans [13] and Neanderthals [14] by producing maximum likelihood co-estimation of sequence error, contamination, and parameters correlated with divergence and heterozygosity. However, this method heavily relies on the significant divergence between archaic and modern humans. A similar method, *DICE*, expanded on this method and jointly estimates contamination rate and error rate along with demographic history based on allele frequency correlation patterns [15]. However, this method requires both explicit demographic modeling and high genome coverage. While this may be effective for estimation of contamination in archaic genomes like Neanderthals and Denisovans that are highly genetically diverged from likely contaminant individuals, it is not optimized for study of contamination among closely related present-day human groups with complex demographic relationships relative to each other, or contamination from individuals of the same population. In Racimo *et al.* 2016 [15], *DICE* required over 3x genome sequence coverage and solved the distinctive problem of measuring contamination of present-day humans in a Neanderthal genome.

We report a method for estimating autosomal aDNA contamination using patterns of linkage disequilibrium (LD) within a sample. This approach, implemented in our software *ContamLD*, is based on the idea that when sequences from one or more contaminating individuals are present in a sample, LD among sequences derived from that sample is expected to be diminished, because the contaminant DNA derives from different haplotypes and therefore should have no LD with the authentic DNA of the ancient individual of interest. Thus, the goal of the algorithm is to determine the LD pattern the ancient individual would have had without contamination and compare it to the LD pattern found in the sample. The LD patterns of ancient individuals are determined using reference panels from 1000 Genomes Project populations to compute approximate background haplotype frequencies, where haplotypes are defined as pairs of SNPs with high correlation to each other. Contamination is then estimated by fitting a maximum likelihood model of a mixture of haplotypes from an uncontaminated individual and a proportion of contamination (to be estimated from the data) from an unrelated individual*. ContamLD* corrects for mismatch of the ancestry of the ancient individual with the reference panels using two different user-specified options. In the first option, mismatch is corrected using estimates from damaged sequences (which, ideally, lack present-day contaminants). In the second option, *ContamLD* performs an “external” correction by subtracting the sample’s contamination estimate from estimates for individuals of the same population believed to have negligible contamination (the user could obtain this value from a *ContamLD* calculation on a male individual with a very low estimate of contamination based on *ANGSD*). The second option has more power than the first option and allows detection of cross-contamination by other ancient samples, but it could be biased if a reliable estimate from an un-contaminated individual from the same population is not available for the external correction.

We show that *ContamLD* accurately infers contamination in both ancient and present-day individuals of widely divergent ancestries with simulated contamination coming from individuals of different ancestries. The contamination estimates are highly correlated with estimates based on X chromosome analysis in ancient samples that are male, as assessed using *ANGSD* [16]. *ContamLD* run with the first option has standard errors less than 1.5% in samples with at least 500,000 sequences covering SNPs (~0.5x coverage for data produced by in-solution enrichment for ~1.2 million SNPs [2, 17], or ~0.1x coverage for data produced using whole-genome shotgun sequences). With the second option, *ContamLD* has standard errors less than 0.5% in these situations, allowing users to detect samples with 5% or more contamination with high confidence so they can be removed from subsequent analyses.

## Results

### Simulations of Contamination in Present-Day Individuals

To test the performance of *ContamLD*, we simulated sequence level genetic data. For our first simulations, each uncontaminated individual was simulated based on genotype calls from a present-day individual from the 1000 Genomes Project dataset. To determine the sequence coverage at each site, we used data from an ancient individual for which we had data at 1.02x coverage and in each case generated the same number of simulated sequences at each site, with the allele drawn from the present-day individual e.g. if the present day individual is homozygous for the reference allele at a site, all simulated alleles are of the reference type, while if the present day individual is heterozygous, simulated alleles are either of the reference or alternative variant, with 50% probability of each). The damage status (i.e. whether it carries the characteristic C-to-T damage often observed in ancient DNA sequences) of each sequence was also determined based on the status of the ancient reference individual. Contaminating sequences were then “spiked-in” at varying proportions (0 to 40%), using an additional present-day individual from the 1000 Genomes Project to determine the contaminating allele type (see Methods). All contaminating sequences were defined to be undamaged, as would be expected if the contamination came from a non-ancient source.

For most of the analyses reported in this study, we simulate data for SNP sites targeted in the 1.24 million SNP capture reagent [2, 17] that intersect with 1000 Genomes sites, after removing sites on the X and Y chromosomes (this leaves ~1.1 million SNPs). The *ContamLD* software also allows users to make panels based on their own SNP sets, and in a later section we report results from a larger panel (~5.6 million SNPs) provided with the software that we recommend for shotgun sequenced samples, and which provides more power to measure contamination. We first analyzed data generated using a reference individual from the 1000 Genomes CEU population (Utah Residents (CEPH) with Northern and Western European Ancestry) and the SNP coverage profile of a 1.02x coverage ancient individual of West Eurasian ancestry (Iberian Bronze individual I3756 who lived 2014-1781 calBCE; see Methods). Supplementary Figure 1 illustrates the distribution of LOD (logarithm of the odds) scores generated when *ContamLD* is run on samples with 0%, 7% and 15% simulated contamination. Supplementary Figure 2 shows the contamination rate estimates generated for data with simulated contamination rates between 0 to 40%. At very high contamination (above 15%) *ContamLD* often overestimates contamination, but in practice samples with above 10% contamination are generally removed from population genetic analyses, so inaccuracies in the estimates at these levels are not a concern in our view (the importance of a contamination estimate in many cases is to flag problematic samples, not to accurately estimate the contamination proportion). *ContamLD* assumes that the individual making up the majority of the sequences is the base individual, so we do not explore contamination rates greater than 50% in these simulation studies.

We observe a linear shift in the contamination estimates such that most estimates are biased to be slightly higher than the actual value, with even greater overestimates occurring at higher contamination rates (Supplementary Figure 2). This is likely due to the difference between the haplotype distribution of the test individual and that of the haplotype panel, since the magnitude of this shift increases as the test individual increases in genetic distance from the haplotype panel. Even in cases where the test individual is of the same ancestry as the haplotype panel (as in Supplementary Figure 2) there is expected to be a shift, because the test individual’s haplotypes are a particular sampling of the population’s haplotypes, and the difference between having only frequencies of the haplotype panel and a particular instantiation of those frequencies in the test individual will lead to the artificial need for an external source (“contaminant”) to fit the model properly.

In contrast to the upward bias in contamination estimates due to mismatch of the individual’s haplotypes with the reference panel haplotype frequencies, we observe negative shifts for inbred individuals, as expected because *ContamLD* assumes the paternal and maternal copy of a chromosome are unrelated. In contrast, if the two chromosomes are related, extra LD will be induced and more contamination will be necessary to produce the expected LD pattern. In principle, this inbreeding effect could be corrected explicitly by estimating the total amount of ROH in each individual and applying this as a correction, although we do not provide such functionality as part of our software. If a reliable methodology for quantifying the proportion of the genome that is affected by inbreeding in ancient individuals becomes available, *ContamLD* could be further improved by using this information as an input parameter.

A final type of bias could be expected to arise if the contamination comes from an individual related to the target individual. In this case the true contamination rate is expected to be under-estimated, because *ContamLD* only detects contamination where the contaminant sequence differs from the target individual’s sequence. If the contaminant carries the same haplotypes as the target individual, in the most extreme case as expected for an identical twin, then the existence of contamination will be missed altogether. In general, contamination from closely related individuals is unlikely to be a concern for many population genetic analyses, as close relatives usually (but with important exceptions) have very similar ancestry.

In our implementation, we correct for these systematic biases in two ways, implemented as different options in *ContamLD*.

The first option leverages information from sequences that contain evidence of the C-to-T damage that is characteristic of ancient sequences. This option assumes these sequences are authentically ancient and not derived from a contaminating source (assumed to be from present-day individuals), so the *ContamLD* estimate based on undamaged sequences is corrected by estimates based on the damaged sequences (see Methods for more details). In the second option, we allow the user to subtract the contamination estimate from the estimate of an individual of the same ancestry assumed to be uncontaminated. An advantage of the second option compared to the first is that it has smaller standard errors (Figure 1), reflecting the fact that it does not rely on estimates from damaged sequences (reliance on damaged sequences reduces power since it often reflects a very small subset of the data). A second advantage of the second option is that it allows estimation of contamination in cases where the source of contamination is also ancient in origin, as would be expected if the contamination occurred anciently or due to cross contamination with other ancient samples (the first option would be expected to produce an underestimate of contamination in such cases, since it assumes that sequences that contain C-to-T damage are not contaminated). On the other hand, a drawback of the second option is that it requires users to identify a relatively high coverage, uncontaminated, ancestry-matched sampled for benchmarking purposes; the method is also only expected to work if there is minimal inbreeding in either the sample of interest or the matched sample. Identifying such benchmarking samples may be impossible when analyzing samples from previously unsampled contexts (e.g. early modern humans), and indeed verifying that a benchmarking sample is uncontaminated is very difficult if it is female (if it is male a method like *ANGSD* can be used).

**Figure 1.**
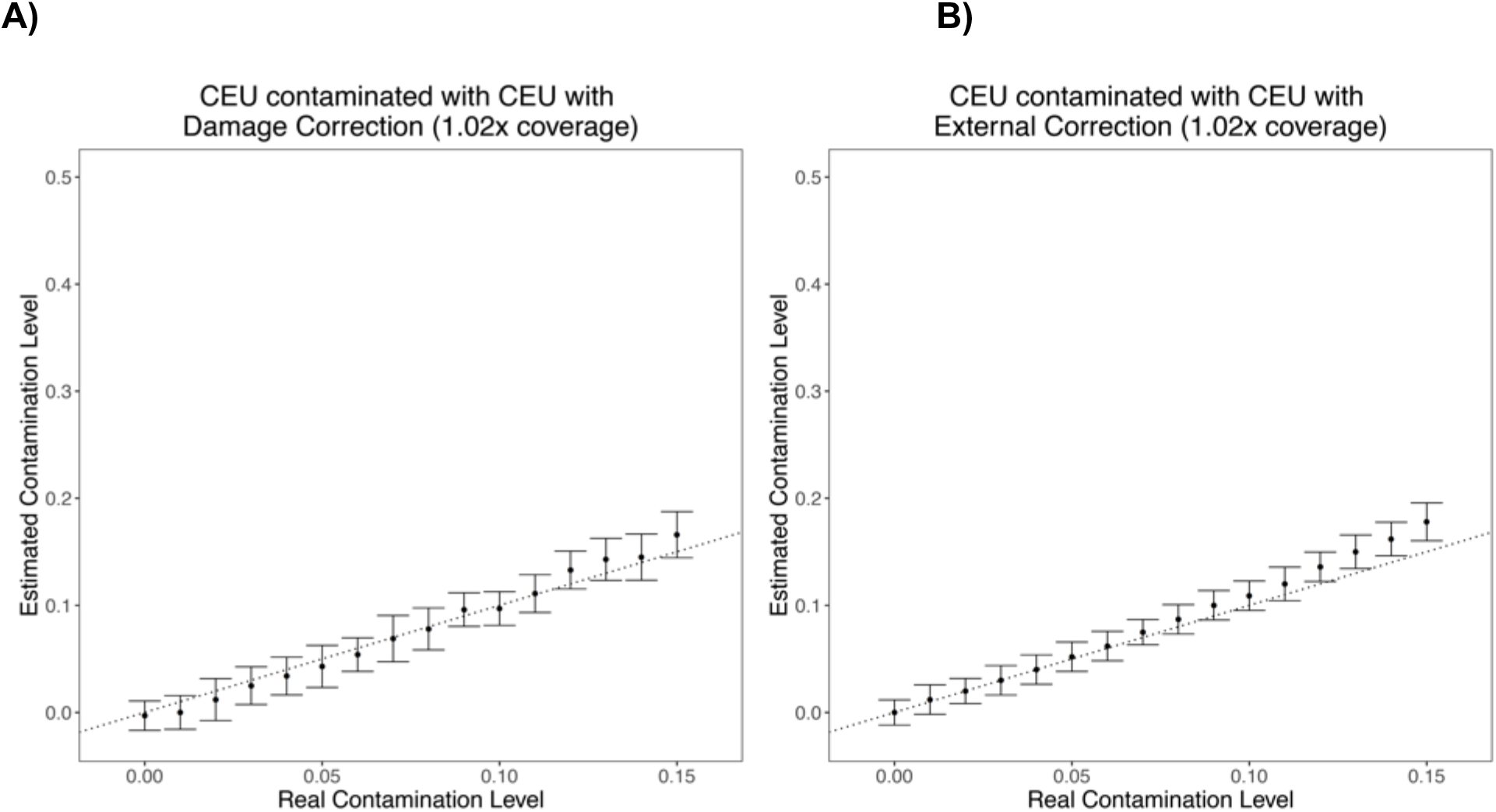
*ContamLD* estimates when the target individual, contaminant, and haplotype panel are from the CEU population. Contamination estimates when the simulated contamination rate is between 0.00-0.15. **A)** Estimates with damage restricted correction (option 1). **B)** Estimates with external correction from an uncontaminated sample (option 2). The black dotted line is y=x, which would correspond to a perfect estimate of contamination. Error bars are 1.96*standard error (95% confidence interval determined via jackknife resampling across chromosomes).

In what follows, we report results of analyses based on the first option, but *ContamLD* includes both methods as options. The uncorrected score also forms the basis for warning output by the software, namely high contamination or possible contamination with another ancient sample leading to an inaccurate damage correction estimate.

### Simulated Contamination of Ancient Samples with Present-Day Samples

*ContamLD* is designed to work on ancient individuals, so we simulated contamination of real ancient individuals with present-day individuals from the 1000 Genomes Project, a scenario that would occur when skeletal material from ancient individuals is contaminated by present-day individuals during excavation or at some point during the processing of the material. We used data from male individuals selected due to very low X chromosome contamination estimates (less than 1%) based on *ANGSD* [16] (developed first in Rasmussen *et al*. [9]; we used method 1 of that software). (We subtracted the *ANGSD* estimates from the *ContamLD* estimates to correct for any residual contamination.) Figure 2A shows results from the Iberian Bronze Age sample [18] (I3756) with 1.02x coverage at the targeted ~1.24 million SNP positions, demonstrating that *ContamLD* produces highly accurate contamination estimates for this simulation.

**Figure 2.**
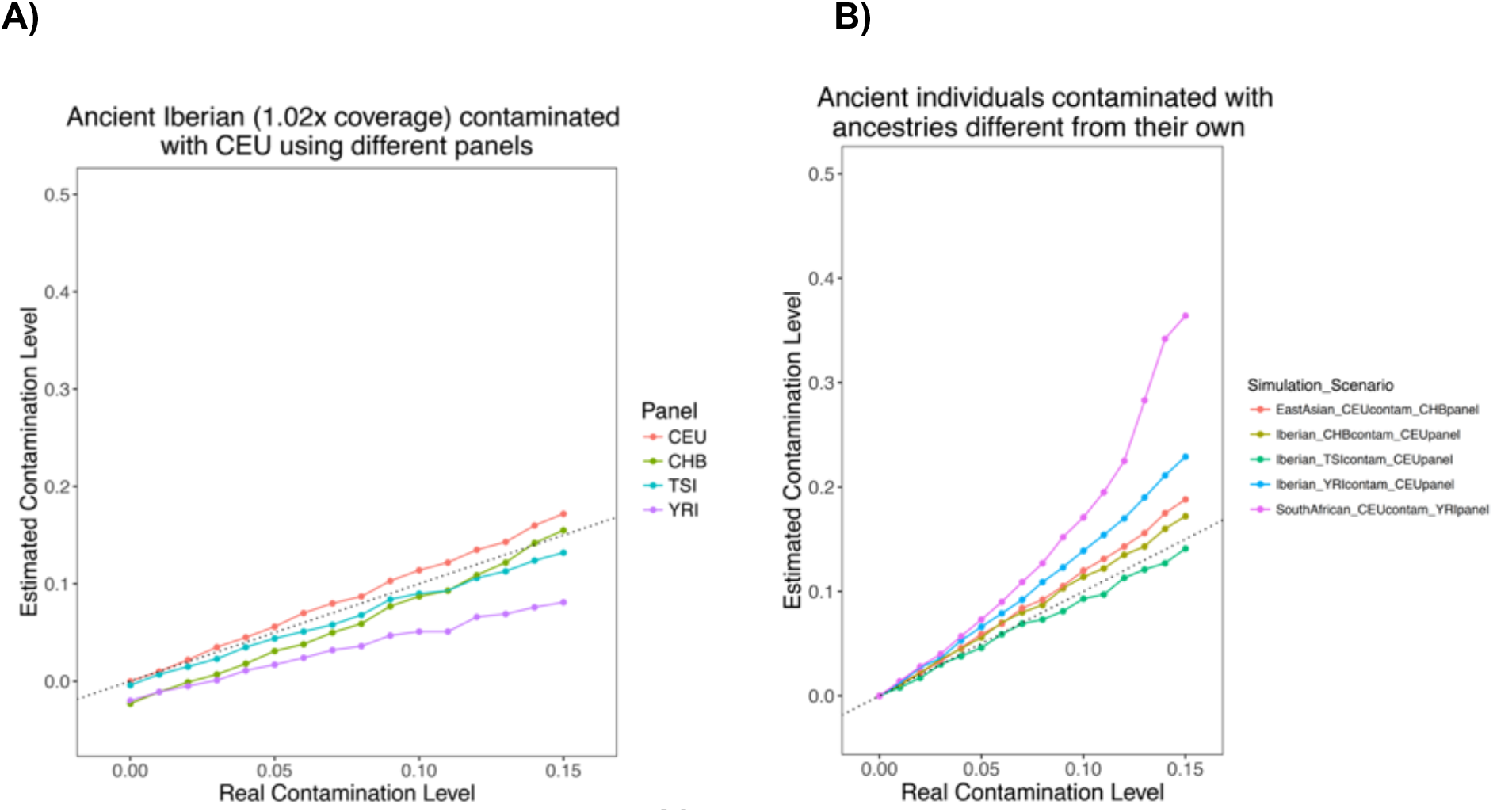
Genetic divergence between uncontaminated individual and contamination sources or haplotype panels impacts *ContamLD* estimates. **A)** Ancient Iberian (I3756, 1.02x coverage) contaminated with CEU with haplotype panels generated from CEU, TSI, CHB, and YRI populations. **B)** Contamination estimates from the same ancient Iberian contaminated with TSI, CHB, or YRI and analyzed with a CEU panel; from an ancient East Asian (DA362.SG, 1.10x coverage) contaminated with CEU and analyzed with a CHB panel; and from an ancient South African (I9028.SG, 1.21x coverage) contaminated with CEU and analyzed with a YRI panel. The black dotted line is y=x, corresponding to a perfect estimate of contamination. All estimates use the damage restricted correction (option 1).

#### Effect of Different Haplotype Panels

There are many potential cases in which ancient individuals can come from populations with very different genetic profiles compared to present-day 1000 Genomes populations, leading to an ancestry mismatch to the haplotype reference panels. *ContamLD* provides panels from all 1000 Genomes populations as well as tools to identify the panel most closely matching to the ancestry of their ancient individual (based on outgroup-*f_3_* statistics [19] to determine the most shared genetic drift), which they can then select for the analysis. However, due to the potential for ancestry mismatch to still occur, we tested the effect of choosing haplotype panels that are genetically diverged from the individual of interest (Figure 2A). For the ancient Iberian sample, the CEU and TSI (Toscani in Italia) panels—representing northern and southern European ancestry, respectively—yielded contamination estimates that are close to the true contamination rate, especially for rates below 5%. However, *ContamLD* underestimates contamination by ~2% when the CHB (Han Chinese in Beijing, China) and YRI (Yoruba in Ibadan, Nigeria) panels were used instead (though we view these as very pessimistic cases, because the user should usually be able to choose a panel more closely related to their ancient individual than these scenarios). We thus recommend that users take care to choose an appropriate panel that is within the same continental ancestry as their ancient individual. Nevertheless, we note that we were able to obtain reasonably accurate estimates for Upper Paleolithic European hunter-gatherers, such as the Kostenki14 individual [20], who is ~37,470 years old, even when using present-day European panels that have significantly different ancestry from the hunter-gatherers (Supplementary Figure 3).

#### Effect of Mismatch Between the Ancestry of the True Sample and Contaminating Individual

Contamination can come from a wide variety of sources, including, but not limited to, members of the archaeological excavation team, the aDNA laboratory, or residual human DNA on the plastic and glassware or in laboratory reagents. Thus, we sought to understand the effect of mismatch in the ancestry of the true sample and the contaminating individual in our contamination estimates. We found that as the ancestry of the two diverged, *ContamLD* over-estimated contamination (Figure 2B and Supplementary Figure 4). This occurred when we tested an ancient European with different contaminant ancestries and when we tested ancient East Asian [21] and ancient South African [22] samples contaminated with European DNA. Nevertheless, the over-estimation was not severe at contamination levels below 5 percent, and samples above this proportion would likely be flagged as problematic. We also explored scenarios where the ancestry of the panel matches the contaminant rather than the true sample (Supplementary Figure 4) and found a ~2% under-estimate at low levels of contamination and an over-estimate at high levels of contamination; these are modest effects and are unlikely to change our qualitative assessment. When we tested the effect of having multiple contaminant individuals (Supplementary Figure 5), we found only a slight over-estimate at higher levels of contamination, as expected given *ContamLD* normally assumes contamination from a single individual where the haplotypes are re-formed if they are created from two contaminant reads (which will happen at lower rates with more contaminant individuals).

#### Estimating Contamination in Admixed Individuals

*ContamLD* relies on measuring the difference between the LD pattern of the sample and that expected from an uncontaminated individual. However, individuals from groups recently admixed between two highly divergent ancestral groups have LD patterns that are similar in some ways to that of an unadmixed individual with contamination from a group with ancestry diverged from that of the individual of interest. To understand how this would impact *ContamLD*, we ran the software on an ASW (Americans of African Ancestry in Southwest USA) individual with different levels of added CEU contamination. When we ran *ContamLD* with a YRI panel and no correction on an individual with no contamination, the individual was inferred to have a contamination of ~20% (likely because the individual had ~15% European ancestry, and this was interpreted by the software as contamination). Using an ASW panel did not perform any better. However, the concerns were mostly addressed by the damage-restricted correction (option 1) at low contamination levels (Supplementary Figure 6). The simulation with African-Americans represents an extreme of difficulty, because the individual is from a group with very recent admixture (~6 generations [23]) of ancestries highly divergent from each other with one of the ancestries very genetically similar to the reference panel. It highlights how the damage-restricted correction is still able to produce accurate estimates in these difficult cases.

### Effect of Coverage

We tested the power of our procedure at different coverages with simulations of ancient West Eurasian ancestry individuals contaminated with CEU on the 1240K SNP set (Figure 3). We found that while our estimates were not biased to produce estimates consistently above or below the true value, the standard errors increased significantly at lower coverages, as expected for the decreased power for accurate estimation in these scenarios. We provide a much larger panel with ~5.6 million SNPs (vs. ~1.1 million for the 1240K panel) that usually decreases standard errors for samples that are shotgun sequenced (Supplementary Figure 7). This panel increases *ContamLD*’s compute time and memory requirements, so we recommend that it only be used for individuals with lower than 0.5x coverage. As an additional feature, we provide users tools to create their own panels to meet their specific needs.

**Figure 3.**
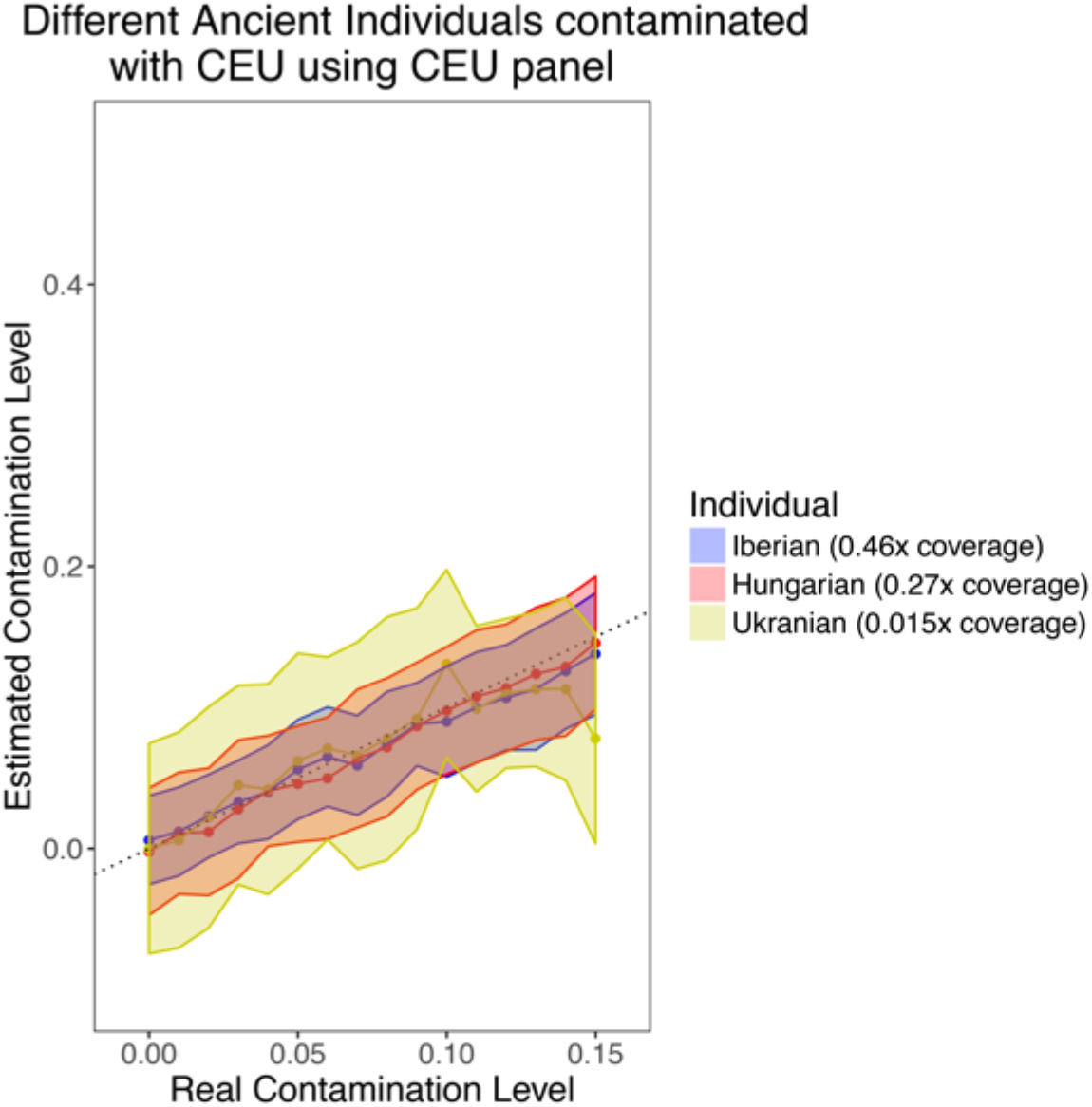
*ContamLD* estimates for ancient European samples of different coverages after damage restricted correction (option 1). An ancient Iberian of 0.46x coverage, an ancient Hungarian of 0.27x coverage, and an ancient Ukranian of 0.015x coverage (~16,000 snps) were contaminated with CEU and analyzed using a CEU panel with *ContamLD* option 1 (damage restricted correction). The black dotted line is y=x. Error shading is 1.96*standard error (95% confidence interval).

#### Simulations to Compare ContamLD to ANGSD X Chromosome Estimates

We performed simulations where we randomly added sequences at increasing levels from 0 to 15% from an ancient West Eurasian individual (I10895) into the BAM files of 65 ancient male individuals of variable ancestries and ages (we set the damaged sequences to be only from the non-contaminant individual; see Methods). We chose ancient male individuals that had average coverage over 0.5X and X chromosome contamination estimates under 2% (using method 1 of *ANGSD*) when no artificial contamination was added (and also corrected even for this baseline contamination by setting damaged reads to be a 5% down-sampling of the files that had no artificial contamination; see Methods). We then analyzed the individuals with *ContamLD* and *ANGSD* and found that compared to *ANGSD*, *ContamLD* consistently had similar errors relative to the true contamination level (Figure 4, Supplementary Online Table 2).

**Figure 4.**
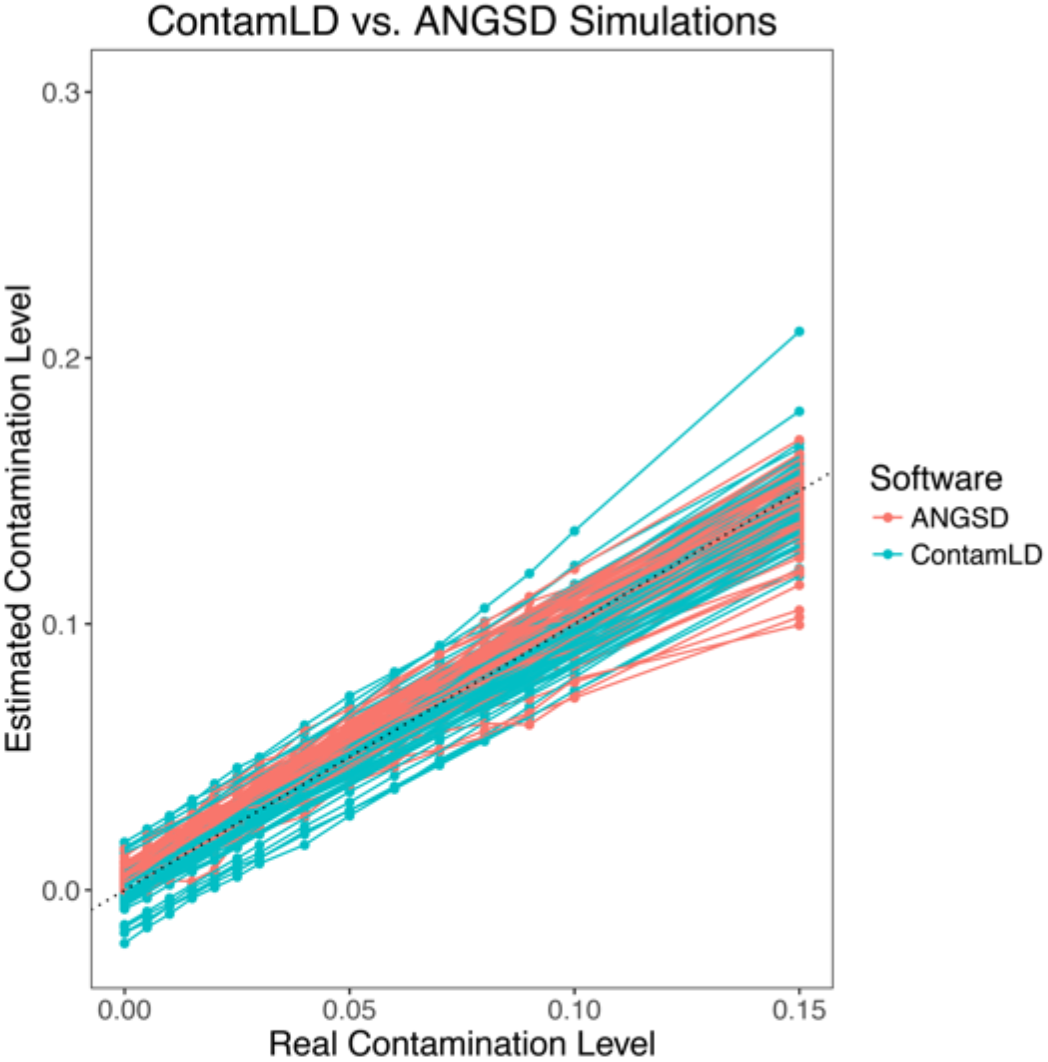
Contamination estimates with *ContamLD* and *ANGSD* for ancient individuals with different levels of contamination added. 65 ancient individuals with average coverage over 0.5X had increasing levels of artificial contamination added in (from I10895, an ~1200BP ancient West Eurasian individual) and were then analyzed with *ContamLD* (with panels most genetically similar to the ancient individual and using damage restricted correction, option 1) and *ANGSD*. Details of all estimates (including standard errors) are provided in Supplementary Online Table 2. The black dotted line is y=x, which would correspond to a perfect estimate of the contamination.

#### Comparing ContamLD, ANGSD, and Mitochondrial Estimates (ContamMix) in Ancient Individuals without Added Contamination

We tested 439 ancient males with *ContamLD*, *ANGSD* (X chromosome contamination estimates), and *ContamMix* (mitochondrial contamination estimates) without adding additional contamination. For this analysis, we included published data generated with the ~1.24 million SNP enrichment reagent, as well as data from libraries that failed quality control due to evidence of contamination (Supplementary Online Table 3). Similar to prior studies [5], the mitochondrial estimates often differed from the nuclear (*ANGSD* and *ContamLD*) estimates, showing high contamination in some libraries with low nuclear contamination, and low mitochondrial contamination in some libraries with high nuclear contamination (Figure 5A). In contrast, *ANGSD* and *ContamLD* had better concordance. However, we observed that some of the samples with high contamination estimates based on *ANGSD* had much lower *ContamLD* estimates, reflecting over-correction from analyzing the damaged sequences, perhaps because the contamination was actually cross-contamination from other ancient individuals, violating the assumptions of our damage-correction (Figure 5B). This problem was mitigated in part, however, because *ContamLD* produces a warning of “Very_High_Contamination” if the uncorrected estimate is above 15% (even in cases where the corrected estimate is very low), and all samples with X chromosome estimates over 5% were flagged with this warning and/or had estimates of over 5% contamination with *ContamLD* (all samples with less than 5% contamination in *ANGSD* had lower than 5% contamination with *ContamLD*). It is unfortunately not possible to know the true contamination of the samples we tested in Figure 5, but the fact that our software produced results with good correlation to X chromosome estimates shows that it works well in real ancient data.

**Figure 5.**
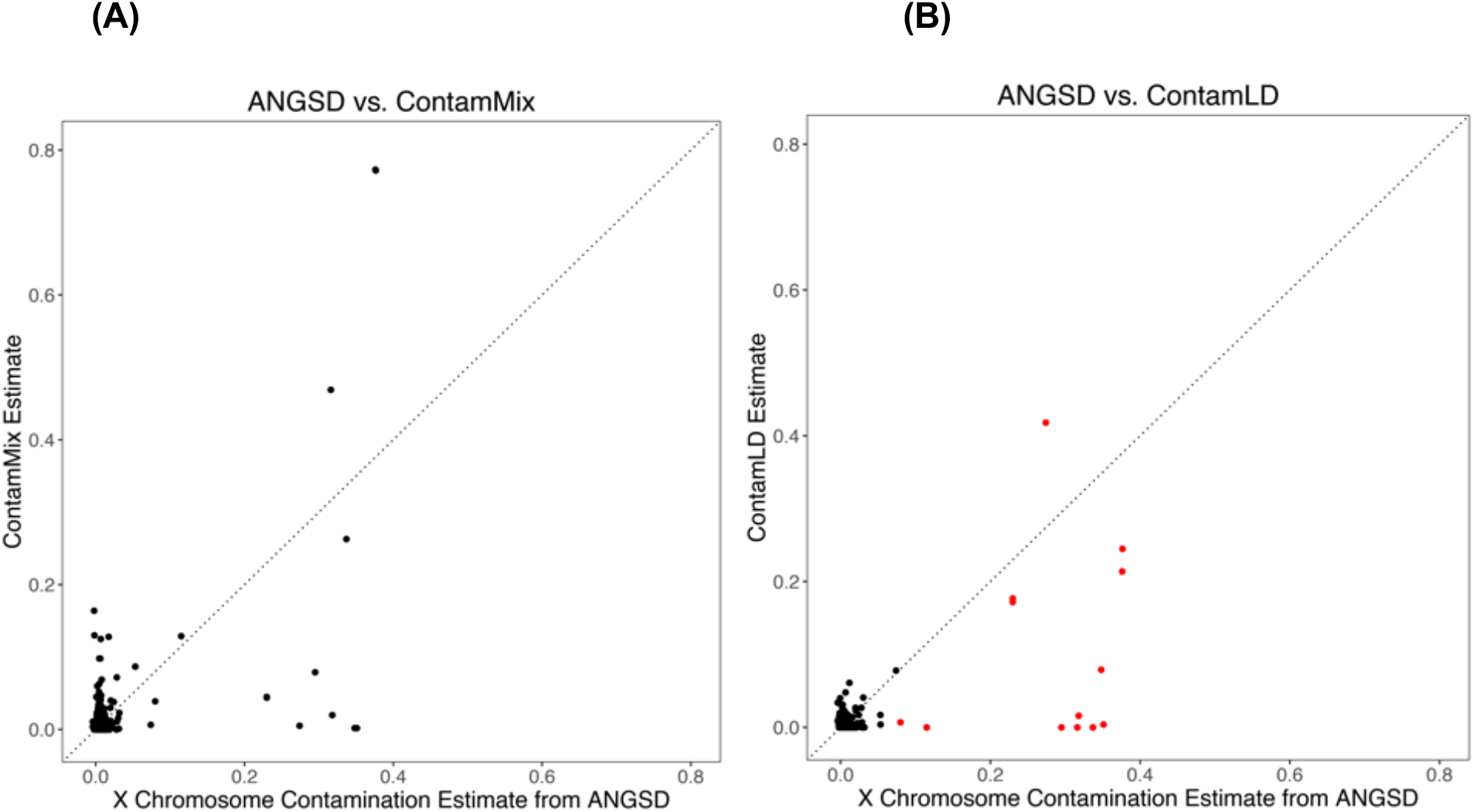
Contamination estimates from *ContamLD*, *ANGSD*, and *ContamMix* in 439 ancient individuals of variable ancestry. *ANGSD* estimates (method 1) are plotted on the X-axis, and on the Y-axis are either **(A)** *ContamMix* or **(B)** *ContamLD* estimates. In red are samples that were flagged in *ContamLD* as “Very_High_Contamination” based on having uncorrected estimates over 15%. All *ContamLD* estimates below 0 were set to 0.

It is possible for there to be samples with moderately high contamination from another ancient individual but both a low damage restricted correction estimate and no warning generated, because these would have high uncorrected estimates, yet not high enough to reach the threshold required for the warning. These samples would have to be identified with an external correction. Lowering the threshold for the “Very_High_Contamination” warning would produce too many false positives, because there are many cases with high uncorrected estimates that have low corrected estimates that are likely not contaminated (e.g. due to ancestry mismatches of the panel and the test individual). To understand these issues better, we performed a simulation in which an ancient Iberian (I3756) was contaminated with another ancient West Eurasian individual (I10895) and the damaged sequences were set to be a 5% down-sampling of the set of contaminated sequences (thus simulating a case in which all of the contamination is from another ancient individual who has the same damage proportion as the ancient individual of interest). We found that, as expected, the contamination from the ancient individual was not detected (the contamination estimates were always near 0%) by the damage restricted correction version of *ContamLD* until the contamination reached 15% at which point the “Very_High_Contamination” flag came up (Supplementary Figure 9). The contamination would have been detected with the external correction version of *ContamLD* (since the damage restricted correction continued to go up with increasing contamination; see Supplementary Online Table 4), but without an uncontaminated ancient individual of the same group as the target individual, this would be difficult to do without bias in the contamination estimate.

## Discussion and Conclusion

We have presented a tool, *ContamLD*, for estimating rates of autosomal DNA contamination in aDNA samples. *ContamLD* is able to measure contamination accurately in both male and female individuals, with standard errors less than 1.5% for individuals with coverage above 0.5X on the 1240K SNP set (for contamination levels less than 10%) for the damage restricted correction method (option 1). On the shotgun panel we provide, standard errors are less than 1.5% for coverages above 0.1x. *ContamLD* is best suited to scenarios in which the contaminant and the ancient individual of interest are similar ancestry, which is useful, because *DICE* [15] and many population genetic tools (e.g. PCA or ADMIXTURE [24]) are better suited for detecting cases where the contaminant is of very different ancestry from the ancient individual of interest. *ContamLD* works even for recently admixed individuals. Lastly, *ContamLD* can detect cases of contamination from other ancient individuals, though this works best if it is large amounts of contamination that can reach the threshold required for the “Very_High_Contamination” flag.

We tested *ContamLD* in multiple simulation scenarios to determine when bias or less reliable results could be expected. When applied to the situation with a test individual (ancient or present-day), contaminant, and haplotype reference panel all from the same continental ancestry, *ContamLD* provides an accurate, unbiased estimate of contamination. When the contaminant comes from a population that is of a different continental ancestry from the population used for the base and haplotype panel, the contamination appears to be slightly overestimated, particularly for higher contamination. This should not be a large problem in analyses of real (i.e. non-simulated) data, because the effect is small at the contamination levels of interest (<5%). When we varied haplotype panels, we found that the estimator is robust when applied to simulated datasets using haplotype panels that are moderately divergent from the base sample (within-continent levels of variation). We provide users tools for automatically determining the panel that shared the most genetic drift with the sample so that the user can select the panel most closely related to the sample. In other simulations, we found that the performance of the algorithm declines as the coverage of the sample decreases. The estimates are not biased, the standard errors substantially increase when fewer than 300,000 sequences are available. In these cases, if the individual was shotgun sequenced, we recommend that users choose the shotgun panel, which will substantially increase power for the analyses.

We applied the algorithm to estimate contamination levels in dozens of ancient samples and compared them to X chromosome-based contamination estimates. There was generally good correlation with the X chromosome estimates, except that when the true contamination was very high, the LD based estimates were sometimes estimated incorrectly, likely because the contamination was due to cross-contamination from another ancient individual and there was over-correction from the damage estimates. This problem is mitigated, however, because the software indicates if the uncorrected estimate is very high so users can identify highly contaminated samples and remove them from further analyses. A difficult case for the software is if there is contamination in part from another ancient sample. This can cause an over-correction and lead to an under-estimate of the contamination. The “Very_High_Contamination” warning catches very high contamination from other ancient samples, but it will miss cases of moderate levels of contamination from other ancient samples, because it will not reach the threshold required for the warning. In theory, the user can determine the true contamination in these cases using the external correction, but the external correction can be difficult if the user does not have an adequate sample to correct the estimate of the sample of interest. The damage correction of the software also does not work if the samples have undergone full UDG treatment (no damaged sequences), and for this case, the external correction is the only option.

The software run-time is dependent on SNP coverage. If ~1,000,000 SNPs are covered (the depth of the coverage on each SNP does not affect run-time), the analysis takes approximately 2 hours if 3 cores are available on CentOS 7.2.15 Linux machines (~25 GB of memory). The software is designed for samples to be run in parallel, so the total time for analysis even for large numbers of samples is often not much greater than the time for a single sample.

In summary, *ContamLD* is able to estimate autosomal nuclear contamination in ancient DNA accurately with standard errors that depend on the coverage of the sample. This will be particularly useful for female samples where X chromosome estimates are not possible. As a general recommendation for users, we believe in most cases all samples with a contamination estimate that is greater than 0.05 (5%) should be removed from further analyses, or the contamination should be explicitly modeled in population genetic analyses.

## Supplementary Data

The Supplementary Data include 4 Excel spreadsheets detailing all ancient samples used and the contamination estimates for this algorithm. Also included are 10 supplementary figures.

## Materials and Methods

### Datasets

#### Present-day samples

Genome wide datasets from individuals that were part of the 1000 Genomes Project [25] were used as present-day reference samples. We restricted to autosomal sites included in the aDNA ~1.24 million SNP capture reagent [2, 17] and to SNPs at greater than 10% minor allele frequency in the pooled 1000 Genomes Project dataset [25]. However, the software allows users to make panels based on their own SNP set. In the analyses presented here, we filtered for SNPs that were present in the 1000 Genomes dataset and also removed all sex chromosome SNPs leading to 1,085,678 SNPs in the final 1240K dataset and 5,633,773 SNPs in the final shotgun dataset.

#### Ancient data set

We analyzed mitochondrial and X chromosome contamination estimates [16, 26] from ancient individuals from previous studies generated by shotgun sequencing or targeted enrichment with 1.24 million SNP enrichment, including many samples that failed quality control due to contamination but were from the same archaeological sites [2, 22, 27–33]. Information about the ancient individual data are detailed in Supplementary Online Table 1 and below.

#### Obtaining sequence information

For each ancient individual, we generated the sequence-depth data from the sample bam file, counting the number of reference and alternative alleles at each SNP site in the analysis dataset. Damage-restricted data was generated by restricting to sequences with PMD scores greater than or equal to 3 [4]. Our software can accommodate both genotype call data as well as sequence data (the sequence data adds additional power to the analyses), but all analyses were performed using the sequence-based method. We provide users with tools to pull down read count data from BAM files in the format required for *ContamLD*.

### Haplotype Calculation

To create haplotype panels, we obtained all SNP pairs in high LD for each 1000 Genomes population using PLINK version 1.9 [34] with r^2^ cut-off of 0.2. (Users can increase power slightly at the expense of increased computational time by creating their own haplotype panel with a lower r^2^ cutoff.) We then calculated the frequencies of each SNP in all of these pairs as well as the haplotype frequencies at each of these pairs while holding out the present-day individuals used for contamination simulation.

### Algorithm to Estimate Contamination

#### Overview

Our goal is to estimate α, the level of contamination, by examining the frequencies of allele pairs that should be in LD (we term this two-allele pair a haplotype) and determining how much their frequencies differ from what would be expected under no contamination. To estimate this, we need both the distribution underlying the haplotypes (*q*) that an uncontaminated test sample should have as well as the distribution of “unrelated haplotypes” (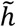) that would form by chance from background allele frequencies. Here and below, “distribution” refers to the set of frequencies of the different possible haplotypes (all possible combinations of ancestral and derived alleles at the SNP pair) across all haplotypes in the genome. Supplementary Figure 10 is a schematic of the algorithm.

#### Determining Haplotype Distributions Based on Reference Panels

To determine *q* we must account for the fact that the test individual’s genotypes do not have diploid calls and are not phased. Due to the low sequence depths at each SNP in many ancient DNA datasets, it is difficult to make confident heterozygous calls, so instead we create pseudo-haploid calls by randomly choosing a sequence to represent the genotype at that position (this holds when we are using genotype calls or the sequence information directly, and when multiple sequences cover the same SNP, we use all of them and treat them as independent). Thus, for this analysis, when examining a pair of SNPs, it is equally likely for the SNP pair to have been formed from the true haplotype (if the same parental chromosome is sampled from in both SNPs of the haplotype) or the background distribution (if the opposite parental chromosome is sampled from). We therefore can estimate *q* as:

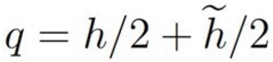

where *h* is the distribution of true haplotypes and 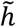 is the distribution of unrelated haplotypes that would form by chance from background allele frequencies. For inbred samples, the weight on *h* is more than 1/2, because the two parental chromosomes are more related, but this can generally be corrected (see below).

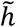 can be determined by multiplying the SNP frequencies to obtain the haplotype frequencies that would form after randomly pairing SNPs of unrelated individuals. *h* can be estimated from an external reference panel using a maximum likelihood estimator (MLE) to obtain haplotypes frequencies in the population from the counts (necessary because the panels are not phased). The MLE set-up is:

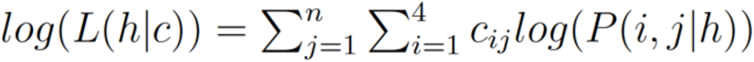

with:

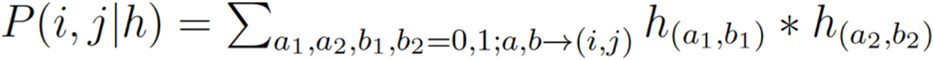

where *P*(*i, j|h*) is the (unknown) diploid count distribution of the haplotypes of the population the test individual is from (approximated by the external panel), *n* is the number of SNP pairs, *c* is the vector of observed haplotypes in the diploid count panel (from 1000 Genomes), i sums over all 4 haplotype possibilities, *h*_(*a,b*)_ are the (also unknown) haplotype distributions of the parents of the test individual (the haploid chromosomes they pass on to their child), and *a, b* → (*i, j*) implies that *a_1_* + *a_2_* = *i* and *b_1_* + *b_2_* = *j*, meaning that one adds up all cases where the haplotype combination would lead to a particular diploid count (e.g. in the notation, for example, 01,11 means the first parent contributes a haplotype that has 0 alternative alleles at the first SNP and 1 alternative allele at the second SNP, and the second parent contributes a haplotype where both SNPs have the alternative allele. The test individual with these parents would then have a 12 diploid count, which means at the first SNP the individual has 1 alternative allele and at the second SNP the individual has 2 alternative alleles. Since our observed data are not phased, both 01,11 and 11,01 would lead to a 12 diploid count). This assumes independence of SNP pairs, which is not true, but because our standard errors are based on jackknife resampling across chromosomes, correlation among SNP pairs is corrected for in our error estimates. The MLE would be computationally intractable to solve due to our lack of knowledge of which parent contributed to each count, so we instead used an EM algorithm to obtain *h*, where knowledge of the parents’ contribution is the unobserved latent variable. The algorithm involves an expectation step of:

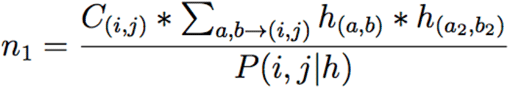

where *n_1_* is the expected number of times that the (*a, b*) configuration of the father’s chromosome contributed to a particular diploid count (this is the same value for the mother, n_2_, because they are assumed to be from the same haplotype distribution). In other words, given the observed haplotype counts in the reference panel, how many times would it be expected that a particular haplotype configuration (e.g. ancestral at SNP1, derived at SNP2) in one of the parents contributed to those counts?

Once the counts (*n_1_* and *n_2_*) of the haploid parents are obtained, they are added together to produce the diploid individual (i.e. the expected number of all possible haplotype configurations). Then the expected value of the haplotype distribution can be maximized by averaging over the possible haplotype distributions. Thus, the maximization step is:

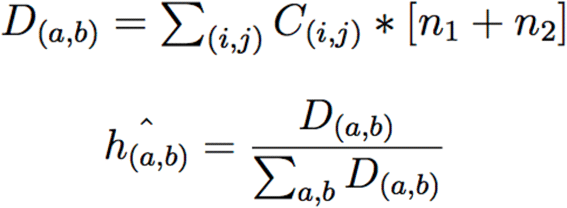

where *D*(*a,b*) is the sum of the probabilities of a particular haplotype configuration over all diploid count configurations.

We initially set all *h*(*a,b*) to be 0.25 and then iterated through the algorithm until convergence (using a squared distance summed over all SNPs and a threshold of 0.001). We then used this estimate of *h* to get an estimate of *q* (based on the first equation above).

#### Estimating Contamination Based on Haplotype Distributions and Test Individual’s Haplotypes

To estimate α, we used the equation:

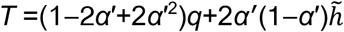

Here *T* is the expected distribution underlying the observed haplotypes of the sample, which is a mix of the test individual and contaminant. This means that assuming the test individual comes from a population with a haplotype distribution (frequency of the different haplotype possibilities at each SNP pair throughout the genome) that can be approximated by the chosen reference panel (and estimated as above), *T* is the haplotype distribution expected for the sample given a particular amount of contamination (*α*’, where ’ is used to indicate that this is an estimate of the real *α*). *q* is the haplotype distribution for an uncontaminated sample. A fraction (1 − *α*’)^2^ + *α*’^2^ of the distribution should look like this, where (1 − *α*’)^2^ is the probability that two uncontaminated sequences form the SNP pair and *α*’^2^ is the probability that two contaminated sequences form the SNP pair, assuming the contaminating sequences are from a single individual, which would “re-form” a SNP pair with LD (note: this also makes the simplifying assumption that the contaminant and the test individual have the same background haplotype and SNP distribution). 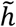 is the distribution of unrelated “haplotypes” that would form by chance from background allele frequencies in the population. Contamination would form these unrelated haplotypes by breaking up LD, so a fraction 2*α*’(1 − *α*’) of the distribution should look like this (the probability that the SNP pair is formed from a contaminated sequence and an uncontaminated sequence).

This expression can be used to solve for α’ by maximizing the LOD (log of the odds) scores under the null hypothesis that *α*’ = 0 and the alternative hypotheses of different *α*’. A LOD score is assigned to each estimate of the contamination rate (*α)* between −0.1 to 0.5 (negative scores are included to allow correction for inbreeding). The grid of *α*’ is scaled by intervals of 0.0001. The *α*’ with the highest LOD score is the best estimate of *α*, and is returned. When we have multiple sequences on the same SNP we assume independence of the sequences, which provides additional power. The assumption of independence does not bias the error estimation for the same reason as explained above for independence of SNP pairs.

#### Correcting for Bias in Contamination Estimates

In practice, the *α*’ we obtain is not equal to the true *α*, because the reference panel does not perfectly capture the SNP and haplotype frequencies of the test sample. We found that this difference causes a linear shift in contamination estimate where the mismatch between the sample individual and the reference panel leads to a positive shift while inbreeding leads to a negative shift. These biases can be addressed in either of two ways.

First, for the “damage correction” approach, we performed an *α*’ estimate only on alleles from sequences with evidence of damage characteristic of ancient samples. Under the assumption that these sequences are not affected by present-day contamination, the inferred *α*’ would be an estimate of the bias, which can be subtracted out from the estimate based on all sites. We separately analyzed the following pairs of SNPs: UU (both SNPs at undamaged sequences), DU (one site damaged and the other undamaged), and DD (both SNPs at damaged sequences). For the UU pairs, the value we calculate would be *α* + k, where k is the linear shift. For DU pairs the value calculated would be *α*/2 + k, and for DD pairs the value calculated would be k. We added the likelihoods for these pairs and maximized the likelihood to solve for *α* and k. After solving for *α*, we multiply by (1-damage rate) to obtain the contamination level across all sequences, because *α* is the contamination rate at undamaged sequences.

Second, for the “external correction” approach, we took individuals from the test individual’s population that were high coverage and samples we believed had very low contamination (based on X chromosome estimates with *ANGSD* using method 1 as developed first by Rasmussen *et al*. [9]) and measured *α*’. We assumed a true contamination of 0 for these samples and thus subtracted this *α*’ from all other contamination estimates. We caution that this method does not correct for uncertainty in the contamination estimate in the external sample used for benchmarking.

#### Comparison to a Similar Method

The approach of *ContamLD* is similar to that of Vohr *et al*. [35] except the two have the opposite goals. Vohr *et al*. searches for LD in reads from two different samples in an attempt to determine whether the two samples are from the same individual (or closely related individuals), using a reference panel to determine LD patterns. In contrast, *ContamLD* searches for breaks in LD in the sequences of a single sample to determine if sequences from other individuals are present in the sample.

### Data simulation

To test the accuracy of the algorithm, we applied it to a variety of scenarios with both present-day DNA as well as real aDNA samples that had simulated present-day DNA contamination. In all our simulations with 1000 Genomes individuals, we removed the individual being used from our haplotype panel before performing the analyses.

#### Simulating Contamination of Present-day Individuals

We first simulated contamination of present-day individuals with other present-day individuals as contaminants (this allowed us to be sure that there was no baseline contamination). In order to best approximate the distribution of both the damaged and undamaged sequences that is characteristic of aDNA data, we used sequence-depth information from an ancient individual as a reference. At each SNP, the total number of simulated “damaged” and “undamaged” sequences was determined based on the number of damaged and undamaged sequences at the SNP in the reference ancient individual. The identity of each allele for the present-day “base” sample was randomly chosen based on the genotype of the “base” present-day 1000 Genomes individual at each SNP, as described above for the contamination. The addition of contaminant sequences to the dataset was performed using the method described above. In order to reduce bias caused by the damage correction procedure, the damage restricted dataset was generated only once for each simulation type (which included multiple simulations across varying contamination rates) and combined with the undamaged dataset to produce the overall dataset. This method was used to generate a simulated individual using present-day CEU (NA06985) or ASW (NA19625) from the 1000 Genomes dataset as the “base” sample from the sequence distributions of a 1.02x coverage ancient Iberian individual (I3756) (the “reference”) [18]. The CEU (NA06984) individual was used as “contaminant” in each case.

We generated simulated data with contamination from multiple sources by adjusting the present-day contamination simulation method to randomly sample from two or more present-day source contaminant genomes with equal probability. In each case, a 1000 Genomes Project CEU individual (NA06985) was used as a “base” genome with the sequence distribution of I3756 (the “reference”). In the case of 2 sources of contamination (Supplementary Figure 5), two CEU individuals from the 1000 Genomes Project dataset (NA06984 and NA06986) were used as contamination sources, and in the case of three contamination sources, an additional CEU individual was used (NA06989). Data was generated for all combinations of undamaged contamination rates, α, from 0-15%.

#### Simulated contamination of ancient individuals

We performed two sets of simulations contaminating different ancient individuals. In both cases we selected ancient male individuals with minimal contamination (as assessed by X chromosome contamination levels from *ANGSD* [16]) to act as the “base” uncontaminated genome. In the first simulation set, we tested *ContamLD*’s performance with different ancient individuals and different present-day contaminant individuals from the 1000 Genomes dataset [25] to assess the impact of contaminant ancestry and coverage of the ancient individual. In this case we were only using *ContamLD* and thus we performed the simulated contamination on the genotype level. In the second simulation set, we compared *ContamLD* to *ANGSD* and used a ~1200BP ancient West Eurasian individual (I10895) to contaminate the BAM files directly.

In the first simulation set, we assumed that sequences with C-to-T damage are highly unlikely to be the product of contamination (this assumption would be falsified in the context of cross-contamination by another ancient DNA sample). Thus, we exclusively added contamination to the “undamaged” fraction of sequences. At each SNP site, we classified sequences present in the damage-restricted dataset as “damaged” and added to the simulated data. We classified all other sequences as “undamaged” and also added them to the simulated data, but for each “undamaged sequence” we added a contaminant sequence to the simulated SNP data with probability *α*/(1-*α*), where *α* is equal to the contamination rate (since the added sequences contribute to the total number of sequences, we needed to add a higher proportion than the contamination rate to obtain our desired contamination rate). The identity of the added contaminant allele was randomly chosen based on the genotype of the chosen “contaminant” present-day genome at the site (i.e. if the contaminant individual was homozygous at the site, the allele it possesses would be added to the simulated individual, while if it were heterozygous at the site, either the reference or alternative allele would be selected randomly and added to the simulated individual). This method maintains the underlying distribution of “uncontaminated” reference and alternative alleles at each SNP site, while adding additional “contaminant” alleles to each site, producing an overall contamination rate of *α* in the undamaged sequences.

For each simulation, we generated two output files: (1) a file reporting the total number of sequences carrying reference and alternative alleles at each SNP and (2) a damage restricted file reporting the total number of damaged sequences carrying reference and alternative alleles at each SNP. We used a 1.02x coverage ancient Iberian individual (I3756) (Supplementary Online Table 1) with contamination from either the 1000 Genomes CEU individual NA06984, the TSI individual NA20502, the CHB individual NA18525, or the YRI individual NA18486. We also used 5 other ancient individuals: I1845 (an ancient Iberian sample of 0.46x coverage) [18], I2743 (an ancient Hungarian of 0.27x coverage) [30], I5891 (a Neolithic Ukrainian individual of 0.016x coverage) [36], DA362.SG (a Russian early Neolithic Shamanka East Asian individual of 1.10x coverage) [21], and I9028.SG (a South African individual of 1.21x coverage) [22]. In each case, we simulated individuals with 0-15% contamination.

For the second simulation set, we analyzed 65 ancient individuals of average coverage over 0.5X and baseline *ANGSD* estimates under 2% (Supplementary Online Table 2). In these cases, we added artificial contamination with sequences from a ~1200BP ancient West Eurasian individual (I10895) into the BAM files at the amounts: (0.000, 0.005, 0.010, 0.020, 0.025, 0.030, 0.040, 0.050, 0.060, 0.070, 0.080, 0.090, 0.100, 0.150). We removed two base pairs from the end of each sequence of partial UDG treated samples and ten nucleotides for non-UDG treated samples and pulled down the genotypes by randomly selecting a single sequence at each site covered by at least one sequence in each individual to represent the individual’s genotype at that position (“pseudo-haploid” genotyping). To ensure that the damage sequences were only from the non-contaminant individual (so that we could use the damage restricted correction mode, option 1, of *ContamLD* without bias), we created the “damaged” sequence set as a randomly chosen 5% of the sequences from the non-contaminant individual. We then analyzed the data with *ContamLD* (damage restricted correction version, option 1) and *ANGSD* using default settings (Method 1). We also performed simulations with a 1.0x coverage ancient West Eurasian ancestry individual (DA57.SG, an ancient Krgyzstanian individual) [37] down-sampled to 0.5x coverage and contaminated with I10895. To simulate different damage rates, we varied the damage rate to the proportions (0.005, 0.01, 0.02, 0.03, 0.04, 0.05, 0.075) by setting the amount of “damaged” sequences to be those proportions.

As a last simulation, we examined the case of an ancient individual contaminating another ancient individual where some of the damaged sequences would also come from the contaminating individual. In this simulation, we analyzed a 1.02x coverage ancient Iberian individual (I3756) and contaminated the BAM with sequences from a ~1200BP ancient West Eurasian individual (I10895) in the proportions (0.000, 0.005, 0.010, 0.020, 0.025, 0.030, 0.040, 0.050, 0.060, 0.070, 0.080, 0.090, 0.100, 0.150, 0.200, 0.300). We then down-sampled the BAM, taking a random 5% of the sequences of these contaminated BAM files to act as the “damaged” sequences, because this would correct for any baseline contamination in the I3756 individual yet would simulate additional contamination of I3756 by an ancient individual with the same damage rate as I3756 (i.e. if there is 5% contamination, then also 5% of the damaged sequences would be from the contaminant individual in this simulation). We then performed the standard processing of both the full contaminated BAMs and the 5% down-sampled BAMs (simulated to be “damaged” sequences), removing two base pairs from the end of each sequence and carrying out a “pseudo-haploid” genotype pulldown. We ran *ContamLD* on the resulting data with damage restricted correction, option 1.

#### Direct Analyses of Contamination Levels in Ancient Individuals

As our last set of analyses, we directly measured contamination levels in ancient individuals without simulated contamination. We used *ContamLD* to examine shotgun sequenced individuals analyzed at the1240K SNP set and the large 5.6 million SNP shotgun panel. The ancient shotgun sequenced individuals were of 0.1-0.5x coverage from Allentoft *et al*., 2015 [31], Damgaard *et al.*, Nature 2018 [37], and Damgaard *et al*., Science 2018 [21]. In addition, we analyzed 439 individuals from a variety of ancestries with *ContamLD* (damage corrected version), *ANGSD* [16, 38] using default settings (we report the results from Method 1), and *contamMix* [39] with the settings: down-sampling to 50X for samples above that coverage, -- trimBases X (2 bases for UDG-half samples and 10 bases for UDG-minus samples), 8 threads, 4 chains, and 2 copies, taking the first one that finishes. Supplementary Online Table 1 includes all information from these individuals.

## Declarations

### Ethics Approval and Consent to Participate

Not applicable (all samples were from previously published studies).

### Consent for publication

Not applicable.

#### Availability of Data and Materials

All data analyzed in this article are available in [2, 21, 22, 27–33, 37]. The software is available at: https://github.com/nathan-nakatsuka/ContamLD. It requires Python 3 and R (any version should suffice). Archived version (1.0) used for analyses in this manuscript: https://zenodo.org/record/3736774#.XoTbj257mgQ (DOI: 10.5281/zenodo.3736774)

Scripts for data simulations are available in the Github folder “Simulation_Scripts”.

#### Funding

Funding was provided by an NIGMS (GM007753) fellowship to NN and a MHAAM fellowship to EH. DR is an Investigator of the Howard Hughes Medical Institute this work was supported by grants HG006399 and GM100233 from the National Institutes of Health, by an Allen Discovery Center grant from the Paul Allen Foundation, and by grant 61220 from the John Templeton Foundation.

#### Authors’ Contributions

N.N., E.H., N.P., and D.R. conceived the study. N.N., E.H., and S.M. performed analysis. N.N., E.H., and D.R., wrote the manuscript with the help of all co-authors.

## Acknowledgements

We thank Iosif Lazaridis and Mark Lipson for helpful discussions.

## Supplementary Figures

**Supplementary Figure 1.**
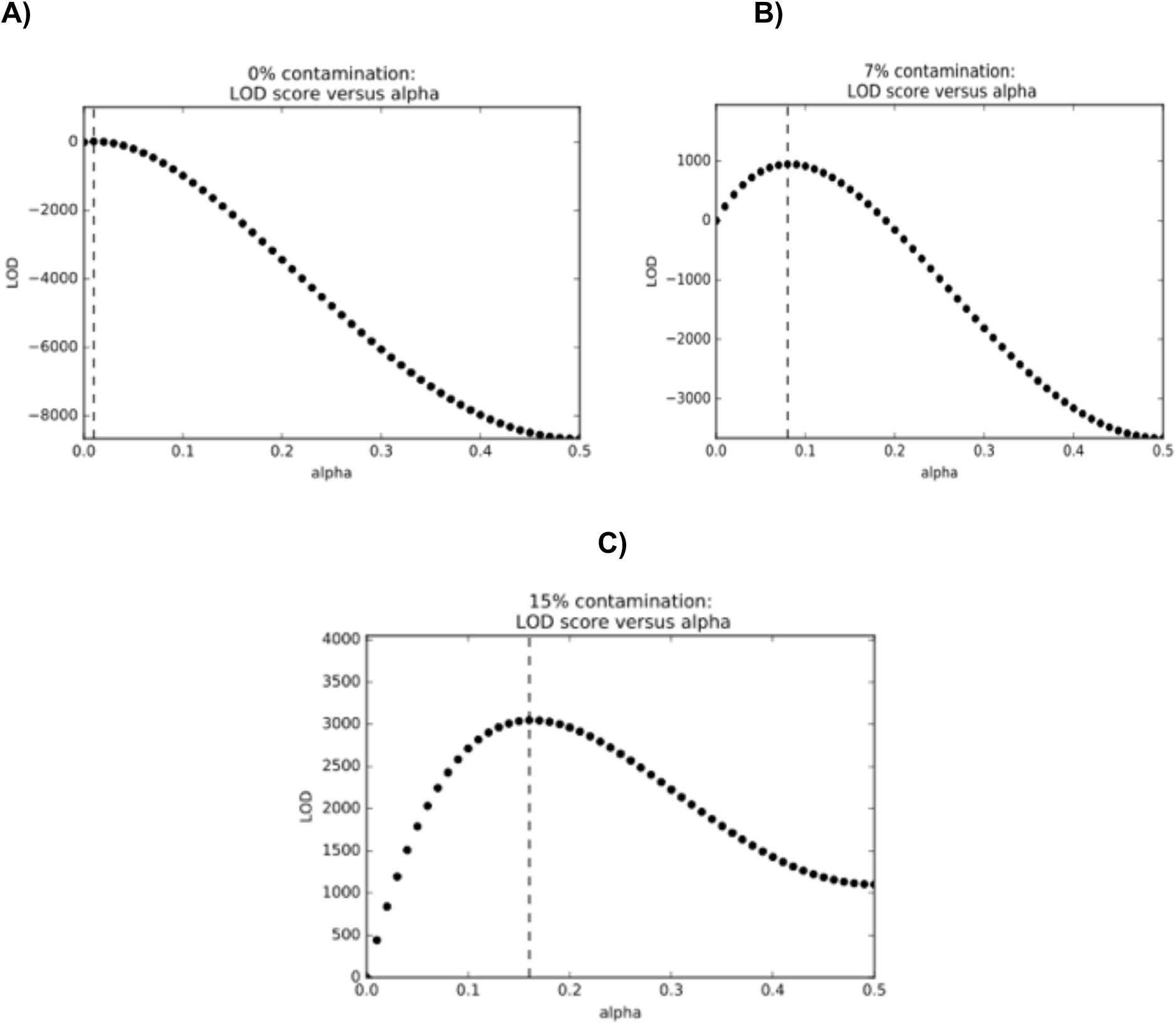
Distribution of LOD scores in simulated data. The distribution of LOD scores is depicted for samples with **A)** 0%, **B)** 7%, and **C)** 15% simulated contamination. These data were generated as part of tests using 1000 Genomes CEU individuals as the sample and contaminant DNA and for the haplotype panel.

**Supplementary Figure 2.**
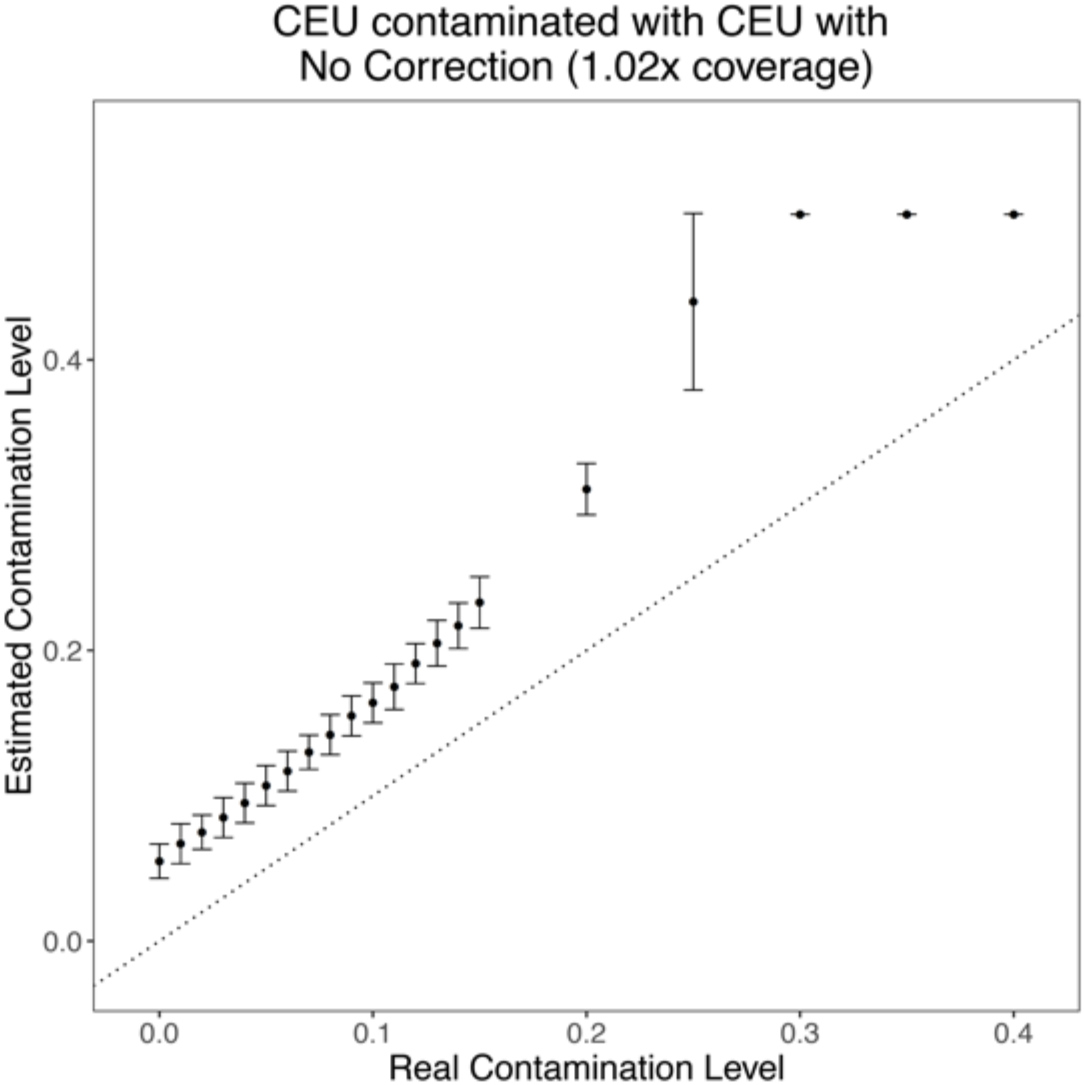
*ContamLD* estimates when the individual, contaminant, and haplotype panel are all from CEU. *ContamLD* was run with no correction. The black dotted line is y=x, which would correspond to a perfect estimation of the contamination. Error bars are 1.96*standard error (95% confidence interval).

**Supplementary Figure 3.**
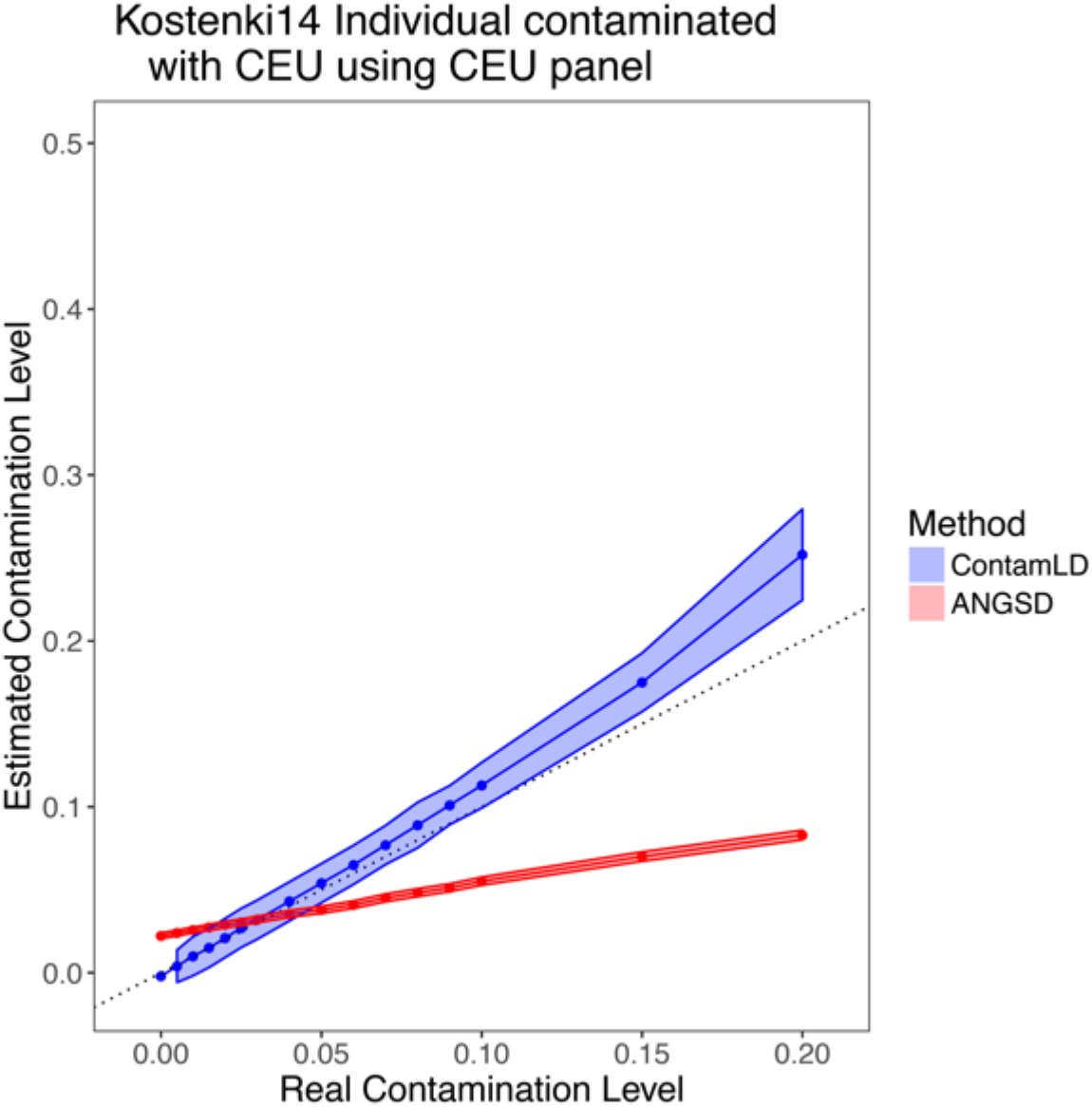
*ContamLD* estimates for Upper Paleolithic European individual after damage restricted correction (option 1). Kostenki14 (2.81x coverage) was contaminated with CEU and analyzed using a CEU panel with *ContamLD* using damage correction and *ANGSD* [16] (Method 1). The black dotted line is y=x, which would correspond to a perfect estimation of the contamination. Error shading is 1.96*standard error (95% confidence interval).

**Supplementary Figure 4.**
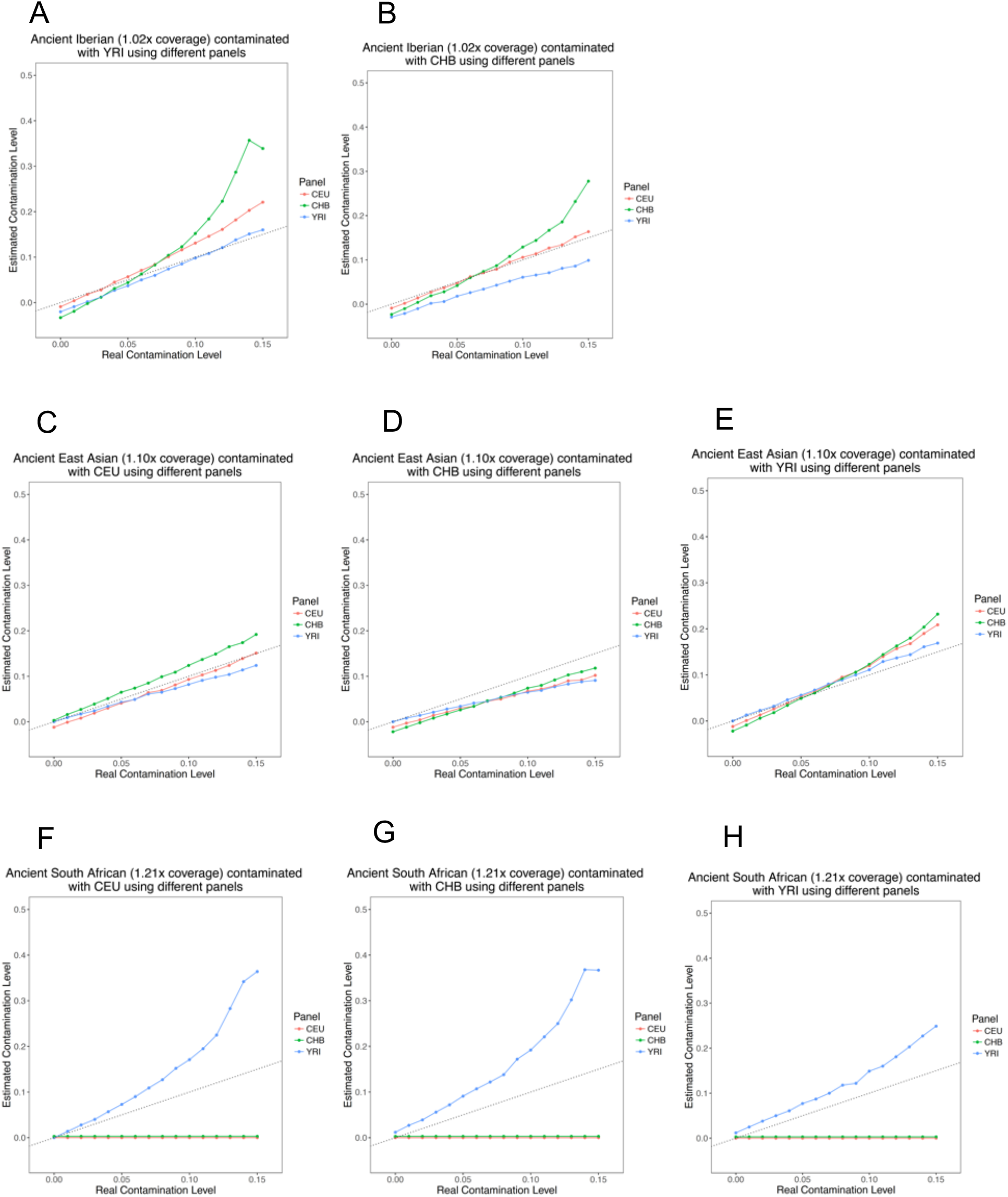
*ContamLD* estimates where the sample, contaminant, and haplotype panels have varying ancestries. *ContamLD* was run with damage restricted correction (option 1). An ancient Iberian of 1.02x coverage (I3756) was analyzed after contamination with **A)** CHB or **B)** YRI. An ancient East Asian of 1.10x coverage (DA362.SG) is analyzed after contamination with **C)** CEU, **D)** CHB, or **E)** YRI. An ancient South African of 1.21x coverage (I9028.SG) is analyzed after contamination with **F)** CEU, **G)** CHB, or **H)** YRI. The black dotted line is y=x, which would correspond to a perfect estimation of the contamination.

**Supplementary Figure 5.**
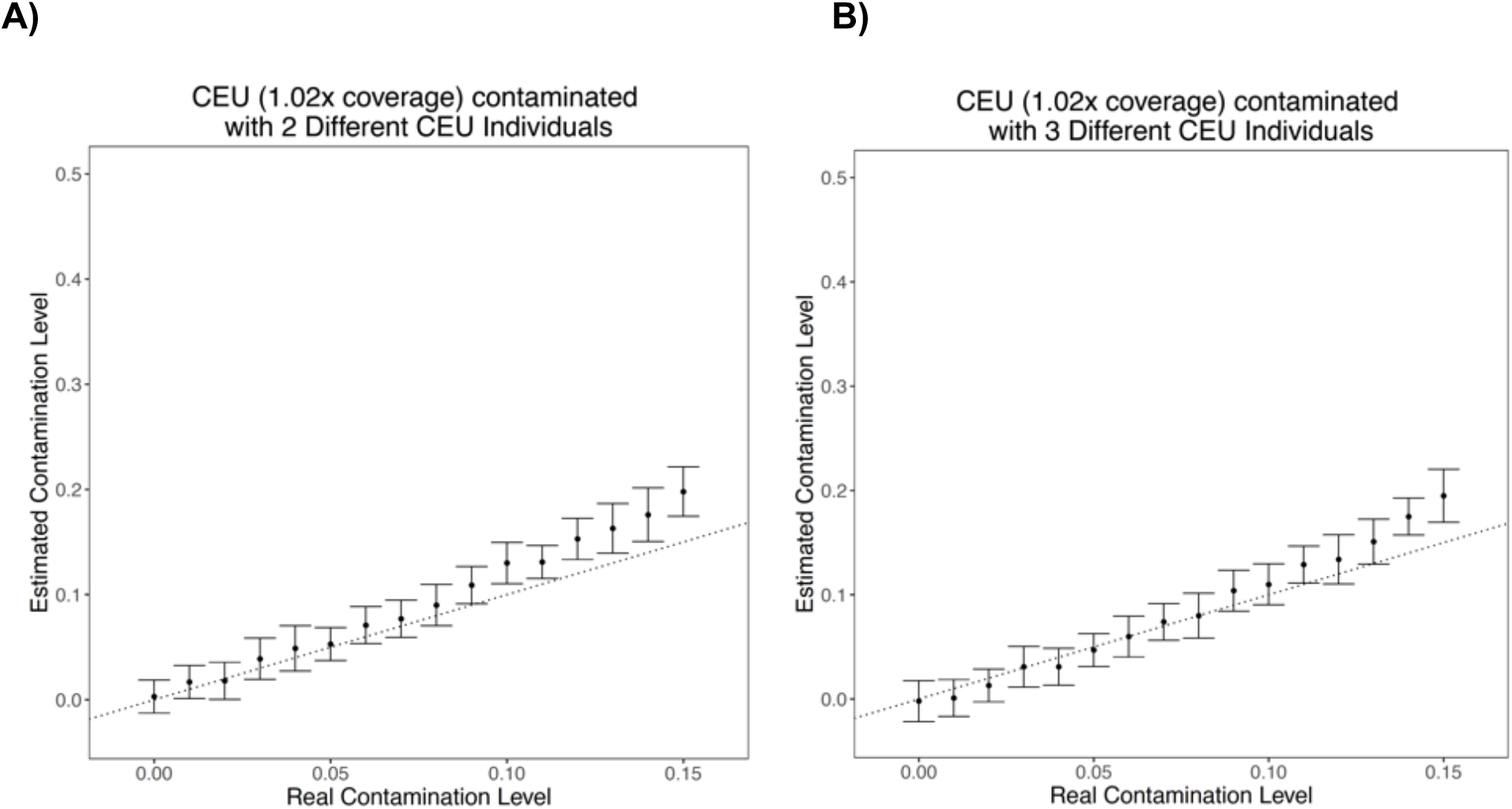
*ContamLD* estimates with CEU as the sample and multiple CEU individuals as contaminants. *ContamLD* was run with CEU haplotype panels and damage restricted correction (option 1). A CEU individual of 1.02x coverage (from the sequence distribution of the ancient Iberian above) is contaminated with **A)** two CEU individuals or **B)** three CEU individuals. The black dotted line is y=x, which would correspond to a perfect estimation of the contamination. Error bars are 1.96*standard error (95% confidence interval).

**Supplementary Figure 6.**
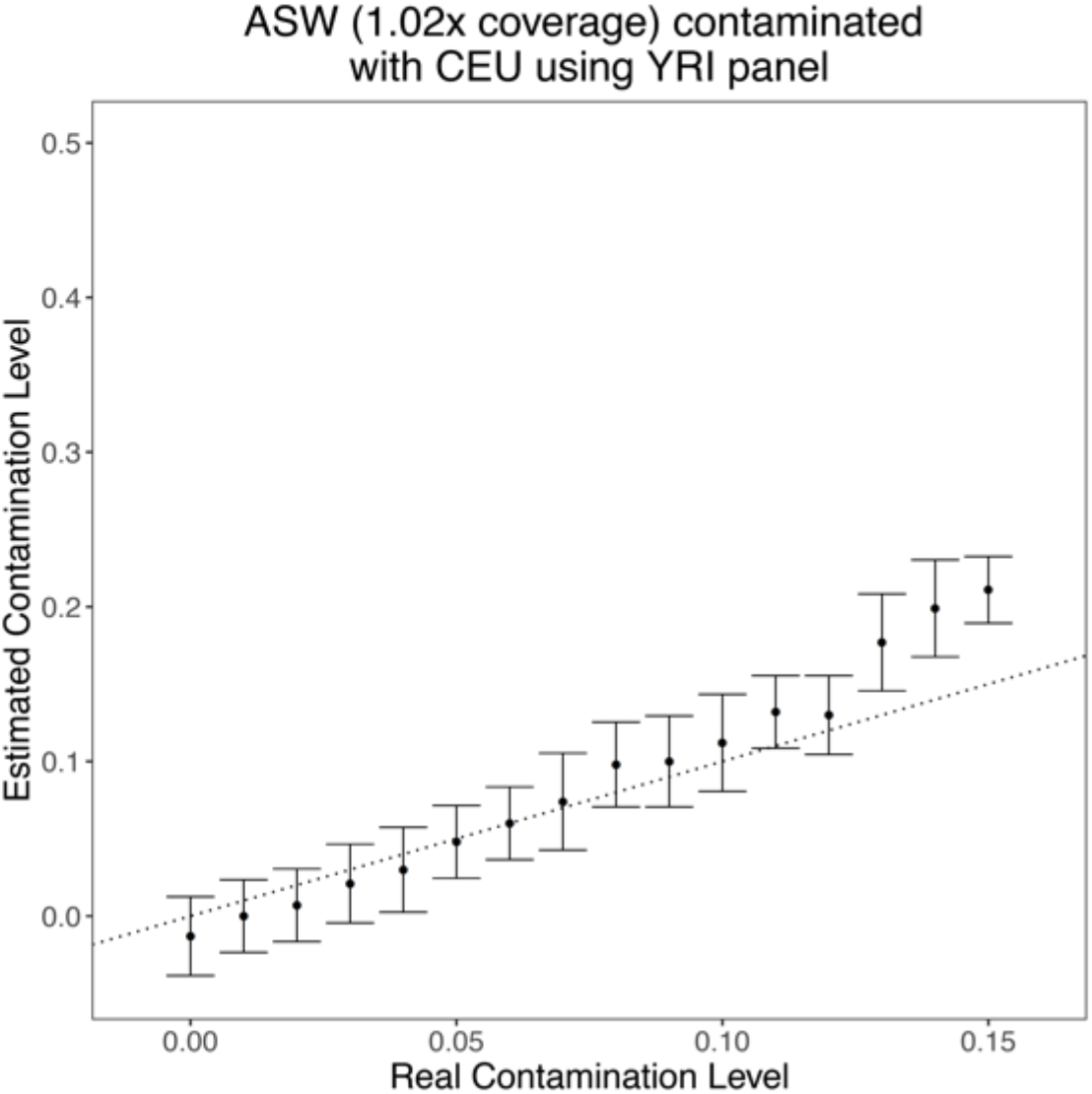
*ContamLD* estimates with an ASW (African-American) individual and YRI panel. *ContamLD* was run with the 1240K SNP set and damage restricted correction (option 1). The black dotted line is y=x, which would correspond to a perfect estimate of the contamination. Error bars are 1.96*standard error (95% confidence interval).

**Supplementary Figure 7.**
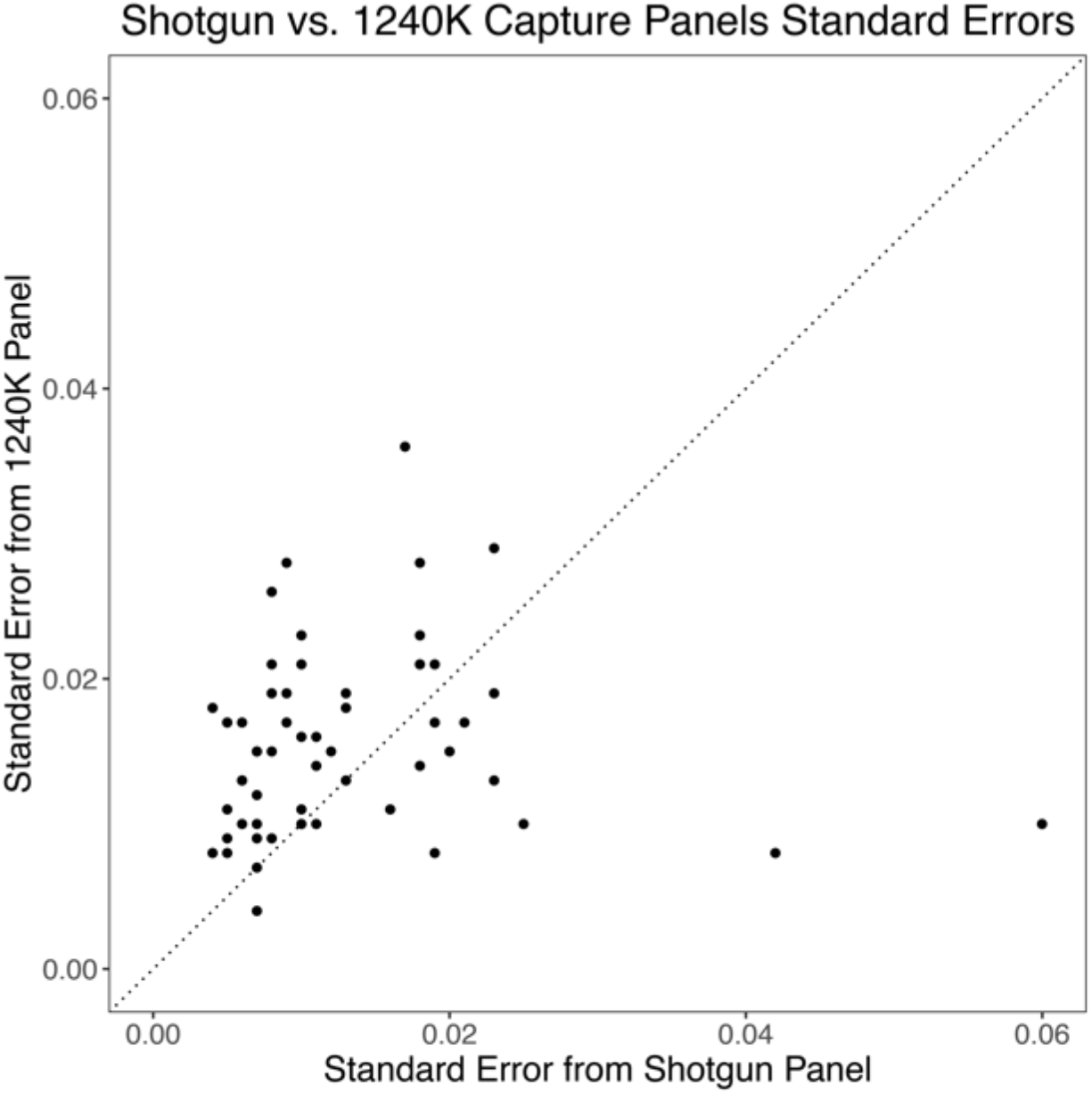
Contamination estimate standard errors of shotgun sequenced ancient individuals comparing the 1240K and shotgun panels. Ancient shotgun sequenced individuals of 0.1-0.5x coverage from Allentoft *et al*., 2015 [31], Damgaard *et al.*, Nature 2018 [37], and Damgaard *et al*., Science 2018 [21] were analyzed with *ContamLD* damage restricted correction (option 1) using the 1240K SNP set and a shotgun panel created using all variants above 10% frequency in the 1000 Genomes dataset. This test shows that analyses with the shotgun panel generally have smaller error bars relative to those done with the 1240K panel, though it is unclear why there are two outliers with high standard errors on the shotgun panel and low standard errors on the 1240K panel. All estimates are in Supplementary Online Table 1.

**Supplementary Figure 8.**
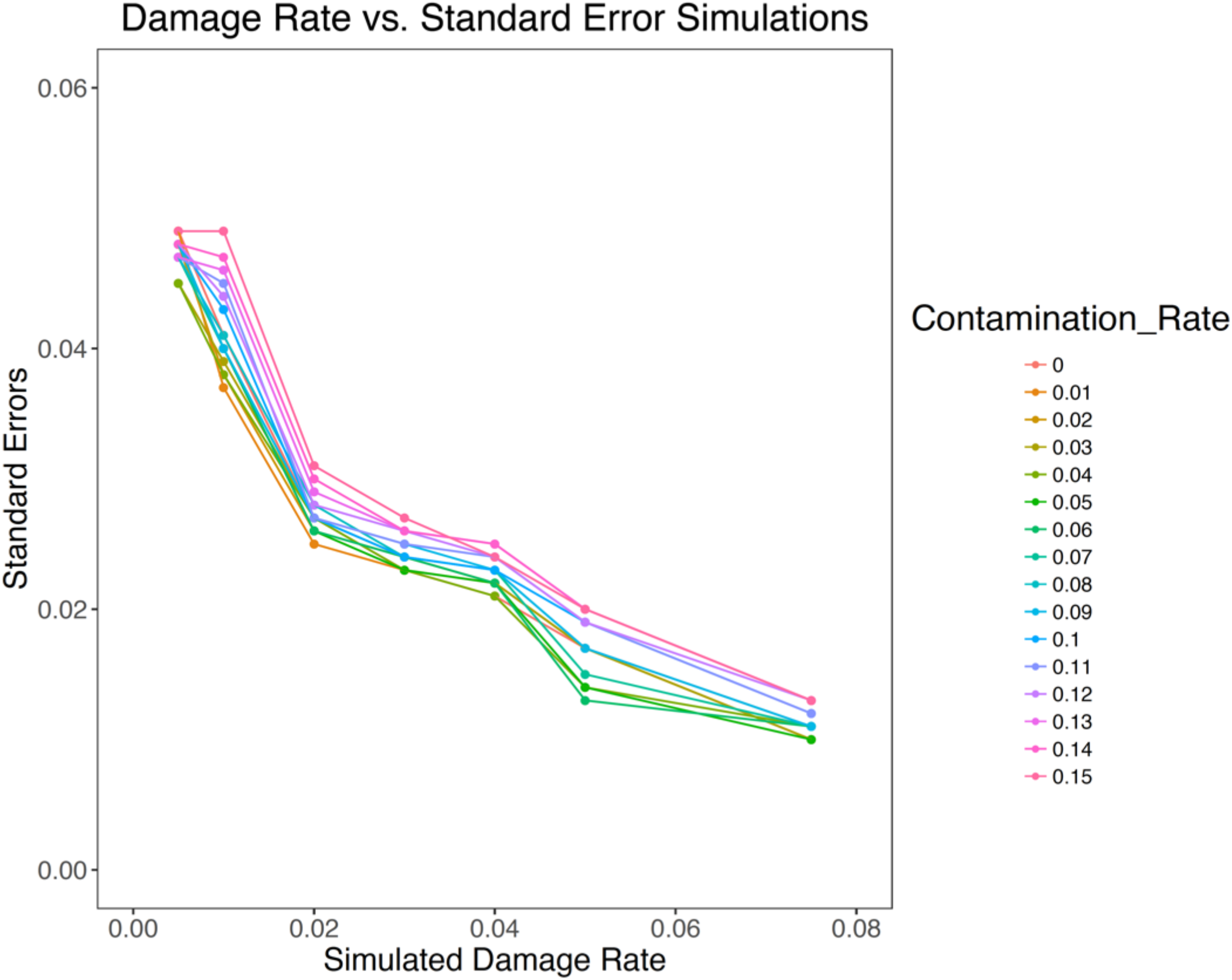
*ContamLD* standard errors from an ancient individual contaminated with another individual at different damage rates. The target individual was a 0.5x coverage ancient West Eurasian-related individual (DA57.SG), and the contaminating individual was an ancient Iberian (I10895). *ContamLD* was run with the 1240K SNP set with the CEU panel. The damaged sequences were simulated as 0.005, 0.01, 0.02, 0.03, 0.04, 0.05, and 0.075.

**Supplementary Figure 9.**
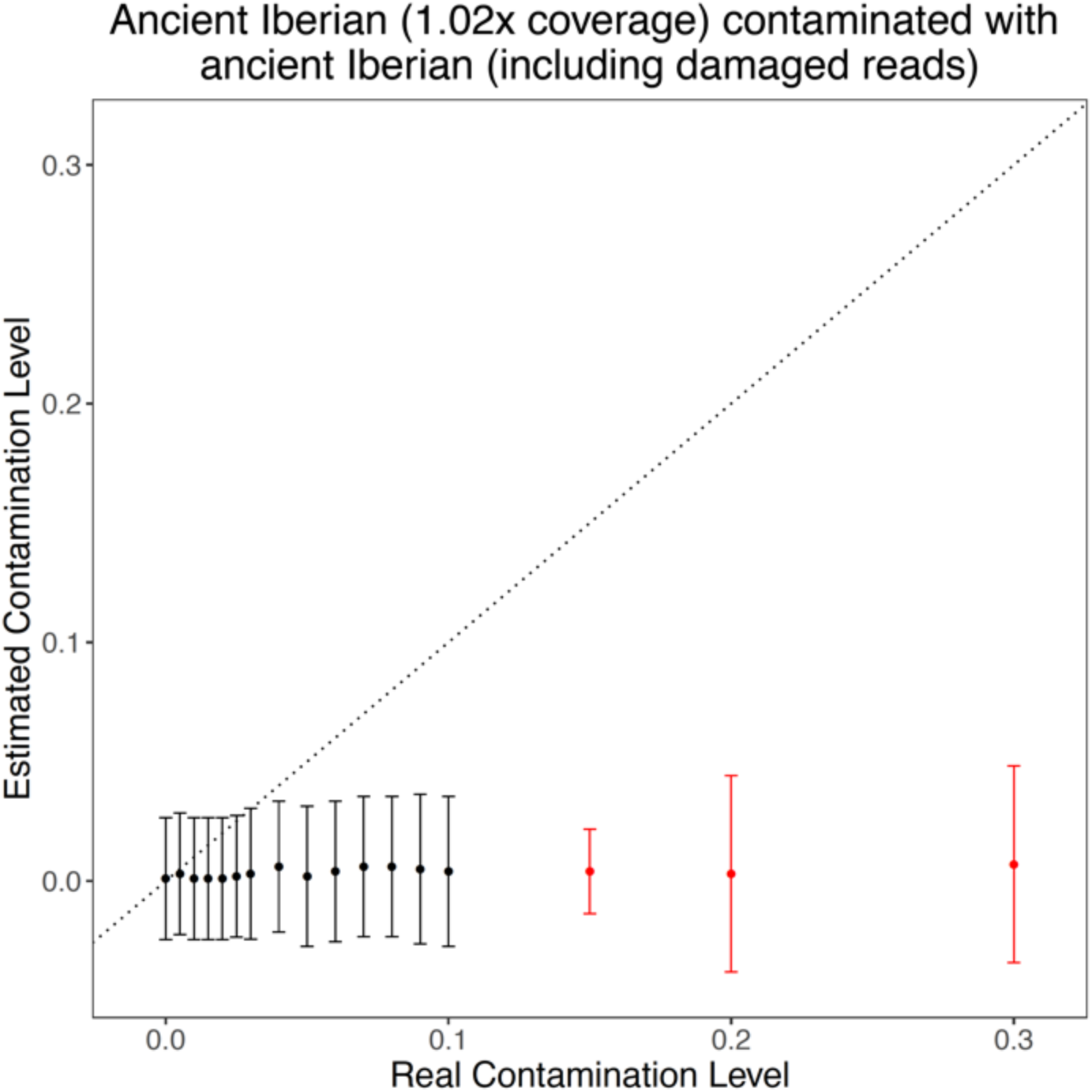
*ContamLD* estimates with an ancient individual contaminated with another ancient individual including its damaged sequences. The target individual was an ancient Iberian (I3756) and the contaminating individual was another ancient Iberian (I10895). *ContamLD* was run with the IBS panel and 1240K SNP set using damage restricted correction (option 1). The damaged sequences were simulated as a 5% down-sampling of each respective contaminated BAM file. IBS are 1000 Genomes Project present-day Iberians from Spain. The black dotted line is y=x, which would correspond to a perfect estimation of the contamination. Error bars are 1.96*standard error (95% confidence interval). Points in red are those flagged with “Very_High_Contamination” by the software. See Supplementary Online Table 4 for all values.

**Supplementary Figure 10.**
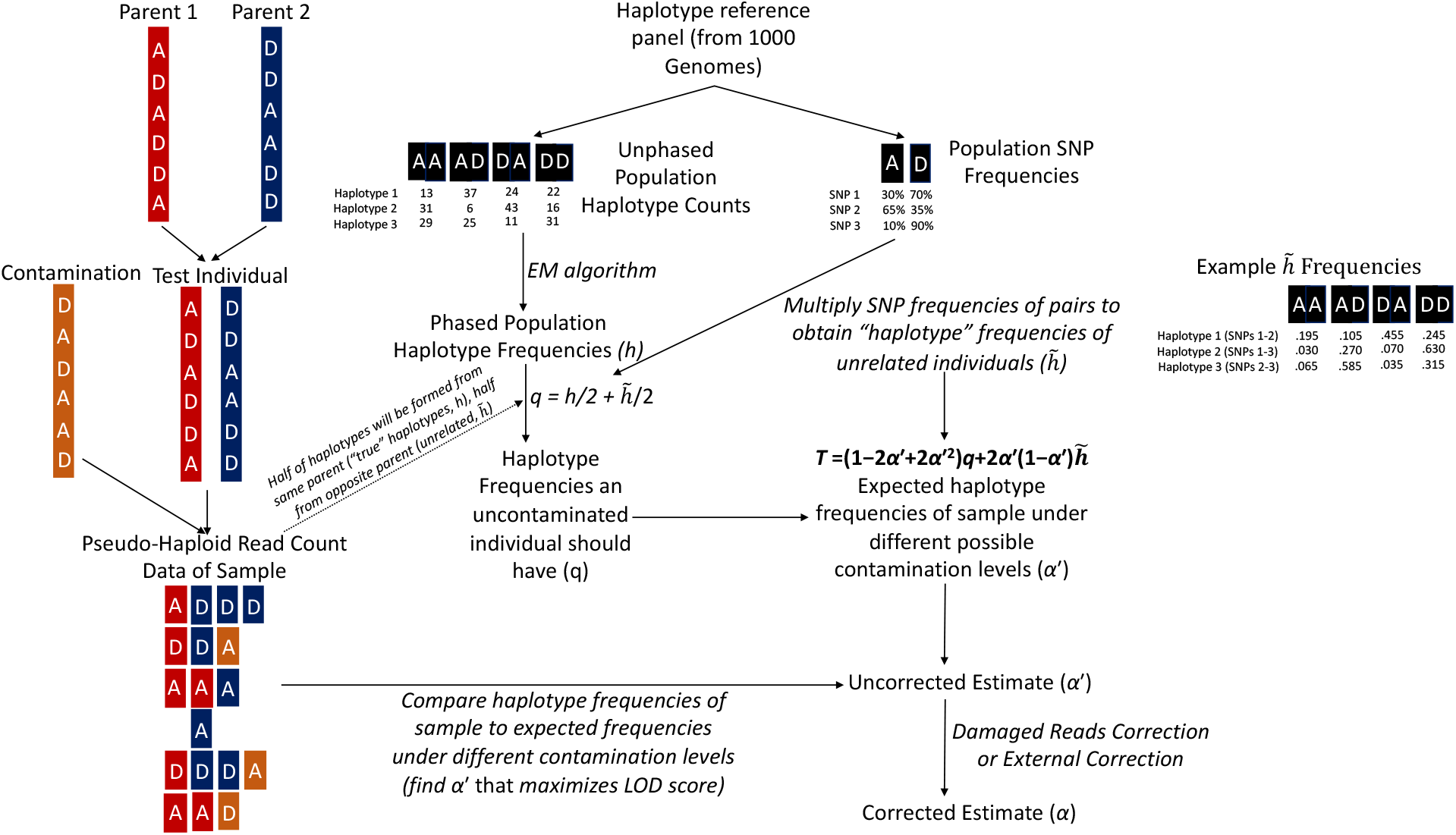
Schematic of *ContamLD* algorithm (see text for additional details).

